# Mature neutrophils promote long-term functional recovery after spinal cord injury in a sex-dependent manner

**DOI:** 10.1101/2025.02.25.640256

**Authors:** Mia R. Pacheco, Ashley V. Tran, Matthew L. Bradley, Miranda E. Leal-Garcia, Mustafa Ozturgut, Ethan A. Barnett, Visruth V. Chakka, Sachit Devaraj, Megan Kirchhoff, Thandiwe Mulamba, Kyndal Thomas, Natalie V. Vu, Dylan A. McCreedy

## Abstract

Neutrophils are abundant and complex innate immune cells that act as first responders to tissue injury, with critical roles in combating infection and initiating would healing. Following spinal cord injury (SCI), neutrophils are the first peripheral immune cells to infiltrate the injured spinal cord in large numbers. Despite the growing body of evidence demonstrating sex differences in neutrophil function, sex as a biological variable in neutrophil responses following SCI has yet to be thoroughly investigated. Here we provide novel evidence that neutrophil responses differ by sex following SCI with divergent effects on long-term outcomes. We show substantial sex-dependent shifts in the phenotype of circulating and intraspinal neutrophils across time following SCI. Depletion of neutrophils immediately after SCI reveals a previously unidentified role for mature neutrophils in promoting long-term functional recovery in a sex-dependent manner. Mechanistically, mature neutrophils acquire an inflammation-resolving phenotype in the acutely injured spinal cord and depletion of mature neutrophils exacerbates long-term macrophage accumulation following SCI in a sex-dependent manner. Collectively, our findings provide the first account of marked sex differences in the response of neutrophils to SCI and elucidate a novel and sex-dependent role for mature neutrophils in promoting resolution of inflammation and long-term recovery following SCI.

## Introduction

Spinal cord injury (SCI) is a devastating and life-altering event marked by irreversible tissue damage and loss of motor, sensory, and autonomic function. The acute phase of SCI is characterized by an extensive inflammatory cascade that substantially influences secondary tissue injury and long-term functional outcomes^1–3^. Neutrophils are the most abundant immune cell (leukocyte) type in human blood and are the first to infiltrate the injured spinal cord in large numbers, peaking within the first day following SCI in rodents^2–5^. While neutrophils have canonically been implicated in collateral tissue damage in the acutely injured spinal cord, recent studies have demonstrated beneficial roles for neutrophils in central nervous system (CNS) injury, including SCI and optic nerve injury^6,7^. Outside of the CNS, there is now a large body of evidence demonstrating that neutrophils play an important role in promoting resolution of inflammation and tissue repair across multiple models of injury, disease, and infection^5,8–19^. The dichotomous role of neutrophils in tissue injury and wound healing may explain the inconsistent effects on long-term outcomes that have been previously reported with neutrophil depletion in murine models of SCI^20–24^. While some of the variability can be attributed to methodological differences, the dynamic and heterogeneous nature of neutrophils, as well as prominent sex differences in neutrophil function^8,25–29^, may also contribute to the discordant effects observed between neutrophil depletion studies. Thus, it is imperative to re-examine the time-, subset-, and sex-dependent roles of neutrophils in recovery processes following SCI.

The multi-faceted nature of neutrophils may be due in part to maturation-dependent changes in neutrophil function. During steady-state conditions, neutrophils predominantly mature within the bone marrow and are released into circulation in a circadian pattern^25,30^. During injury or infection, however, immature neutrophils are promptly released from the bone marrow in large numbers resulting in a rapid increase in circulating immature and mature neutrophils (neutrophilia). Recent studies support neutrophil maturation as a basis for phenotypic diversity in neutrophils, and disparate roles for mature and immature neutrophils have been reported for tumor progression^25,26,31^, lymphocyte regulation^32^, axon regeneration^28^, and pathogen clearance^33^. However, neutrophil heterogeneity and maturation in the context of SCI, have yet to be characterized.

Prominent sex differences in neutrophil phenotype and functional responses have been recently reported for mice and humans. In humans, sex differences have been observed in neutrophil extracellular trap (NET) formation, reactive oxygen species (ROS) production, apoptosis, metabolism, transcriptome, chemorepulsion, and the maturation profile of circulating neutrophils^27,34–37^. In mice, multi-omic profiling of primary bone marrow neutrophils has demonstrated sex-dependent transcriptional signatures and chromatin accessibility profiles^38,39^. Despite the recent emphasis of sex as a biological variable in SCI research, sex differences in neutrophil responses to SCI, as well as the sex-dependent contribution of neutrophils to post-SCI recovery, remain unclear.

In this study, we provide novel evidence of sex differences in granulopoiesis and neutrophil mobilization following injury in a clinically relevant murine model of contusive SCI. Our findings demonstrate, for the first time, that the phenotype of circulating and intraspinal neutrophils rapidly and robustly changes in a sex-dependent manner after SCI. In addition, we show that neutrophils mediate long-term functional recovery following SCI in a sex- and time-dependent manner. Mechanistically, we show a novel reparative role for mature neutrophils in the injured spinal cord, and provide evidence that neutrophils regulate resolution of inflammation in a sex-dependent manner following contusive SCI. Collectively, our data provide critical insight into how neutrophil maturation, temporal responses, and sex differences contribute to long-term outcomes after SCI.

## Results

### Systemic neutrophil responses differ by sex after SCI

To characterize sex- and time-dependent changes in systemic neutrophil responses to SCI, we performed a moderate thoracic contusion SCI in adult male and female mice followed by flow cytometry analysis of blood and bone marrow samples collected up to 35 days post-injury (dpi) (Fig. 1a-b). We observed a rapid increase in circulating neutrophils (neutrophilia) by 4 hours post-injury (hpi) with sustained neutrophilia for at least 1 day following SCI in both sexes (Fig 1c). Interestingly, male mice had a substantially greater neutrophil mobilization response compared to females, with neutrophilia persisting for at least 1 week following SCI. While we observed injury-induced neutrophilia in female mice, the abundance of circulating neutrophils returned to pre-injury baseline by 3 dpi. We next characterized the maturation phenotype of circulating neutrophils in male and female mice. In both sexes, there was marked reduction in the relative proportion of mature (CD101^+^) neutrophils early after SCI with neutrophils displaying a predominantly immature (CD101^−^) phenotype by 1 dpi (Fig 1d-e). In addition to having a larger neutrophil mobilization response post-SCI, male mice also had a greater proportion of mature (CD101^+^) circulating neutrophils than female mice after SCI. Neutrophil maturation profiles recovered to pre-injury levels by 3 dpi in male mice, whereas neutrophils in female mice did not return to pre-injury levels until after 3 dpi.

**Figure 1:**
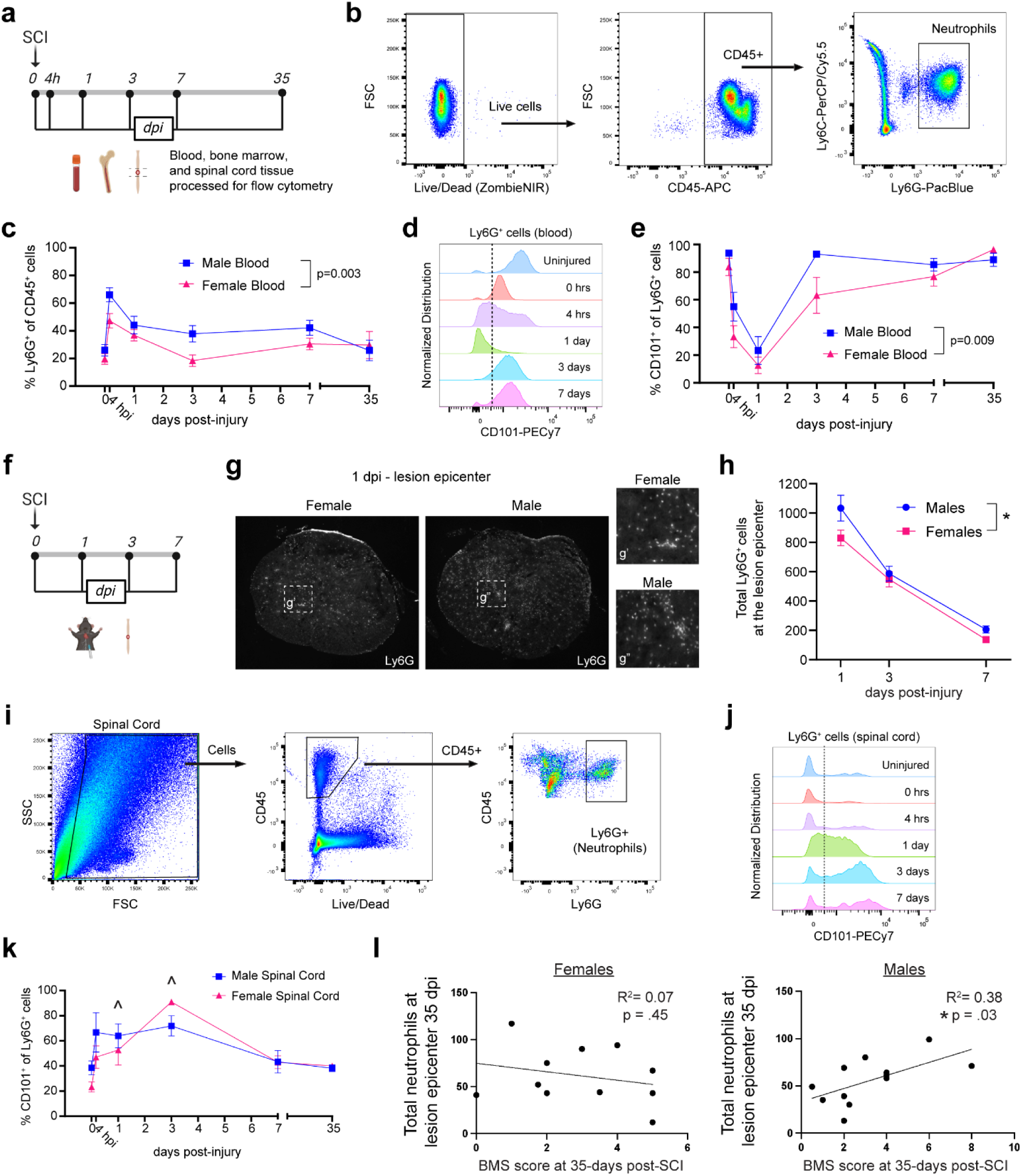
Neutrophil responses differ by sex after spinal cord injury. (a) Schematic of SCI and tissue collection timepoints for flow cytometry analysis. (b) Representative flow cytometry gating of Ly6G^+^ neutrophils in blood samples. (c) Quantification of neutrophil (Ly6G^+^) percentage of total circulating leukocytes (CD45^+^) after SCI. Mixed effects analysis with Sidak’s multiple comparisons test. n=3-10/sex/timepoint. (d) Representative histograms for CD101 levels on circulating neutrophils (Ly6G^+^ cells). (e) Quantification of the percentage of mature neutrophils (CD101^+^/Ly6G^+^ cells) out of total neutrophils (Ly6G^+^ cells) via flow cytometry in the blood after SCI. Mixed effects analysis with Sidak’s multiple comparisons test. n=3-5/sex/timepoint. (f) Schematic of SCI and spinal cord tissue collection for histology. (g) Representative images of Ly6G immunostaining (neutrophils) in the lesion epicenter at 1 dpi in female (left) and male (right) mice. g’ and g’’ are magnified insets. (h) Quantification of Ly6G^+^ cells in the lesion epicenter at 1, 3, and 7 dpi. Mixed effects analysis with Sidak’s multiple comparisons test. n=8-11/sex/timepoint. *p<0.05. (i) Representative flow cytometry gating of neutrophils (Ly6G^+^ cells) in spinal cord samples. (j) Representative histograms for CD101 levels on intraspinal neutrophils (Ly6G^+^ cells). (k) Quantification of the percentage mature neutrophils (CD101^+^/Ly6G^+^ cells) out of total neutrophils (Ly6G^+^ cells) via flow cytometry in the spinal cord after SCI. Mixed effects analysis with Sidak’s multiple comparisons test. n=3-6/sex/timepoint. ^p<0.05 compared to 0 hpi. (l) Correlation of BMS scores and intraspinal neutrophils at the lesion epicenter in female (left) and male (right) mice at 35 dpi. Linear regression analysis. n=11/sex. Abbreviations: SCI = spinal cord injury, BMS = Basso Mouse Scale, dpi = days post-injury. Mean +/− SEM.

We also characterized the effect of SCI on bone marrow neutrophil populations. As expected, there was a reduction in the proportion of mature neutrophils in the bone marrow in both male and female mice at 1 dpi, likely due to the generation of new immature neutrophils as well as mobilization of reserve mature bone marrow neutrophils into circulation early after injury (Fig S1a). At 3 dpi, we observed a sex-dependent recovery of the bone marrow neutrophil populations, which returned to pre-injury mature neutrophil levels (∼20%) in male, but not female mice. EdU labeling of proliferative neutrophil precursors 1 day prior to SCI revealed that females had a higher proportion of newly generated immature neutrophils (CD101^−^/EdU^+^) in the bone marrow at 1 dpi (Fig S1b-d), whereas males had a higher proportion of new mature neutrophils in the bone marrow at 3 dpi (Fig S1e). Consistent with the increase in the abundance of mature neutrophils in the blood of male mice seen at 3 dpi, there was a greater percentage of newly generated mature neutrophils (CD101^+^/EdU^+^) in the blood of male mice relative to females at 3 dpi (Fig S1f). Collectively, our data provide novel evidence of sex differences in SCI-induced neutrophilia, including sex-dependent changes in neutrophil maturation and granulopoiesis in the bone marrow and blood after SCI.

### Intraspinal neutrophil accumulation and phenotype differ by sex after SCI

Since we observed prominent sex differences in neutrophilia and the maturation profile of circulating neutrophils following SCI, we also assessed if there were sex differences in neutrophil accumulation and phenotype in the injured spinal cord. In accordance with our findings that male mice experience greater neutrophilia after SCI, we observed higher neutrophil counts at the lesion epicenter across the first week post-SCI in male mice relative to females (Fig. 1f-h). Flow cytometry analysis of neutrophil maturation in the injured spinal cord revealed a marked increase in mature neutrophils in the injured spinal cords of female mice at 1 and 3 dpi relative to 0 hpi (Fig. 1i-k). While there was a modest trend towards an increase in the mature neutrophil proportion in male mice at 3 dpi compared to 0 hpi, the percentage of mature neutrophils in the spinal cord at 3 dpi was greater in female mice relative to males (Fig. S1g). Altogether, we observed that the accumulation and maturation phenotype of intraspinal neutrophils rapidly changes in a sex-dependent manner after SCI.

To assess if neutrophil lifespan is extended in the injured spinal cord, we assessed EdU persistence in intraspinal neutrophils in mice pulsed with EdU at 1 day prior to SCI. While EdU was no longer detected in neutrophils in the bone marrow and blood by 7 dpi (Fig. S1e-f), we still observed EdU^+^ neutrophils in the injured spinal cord up to 35 dpi in both sexes (Fig. S1h-i). We also assessed if neutrophils can enter the spinal cord in the subacute phase of SCI by pulsing mice with EdU at 3 weeks post-SCI. We detected newly generated neutrophils (EdU^+^) in the injured spinal cords of both male and female mice at 35 dpi (Fig. S1j-k), confirming that neutrophils can enter the spinal cord, albeit in limited numbers, weeks after injury. To the best of our knowledge, our data are the first to demonstrate extended neutrophil lifespan after SCI, as well as continued infiltration of the injured spinal cord by newly born neutrophils.

We next examined whether there is a relationship between chronic neutrophil accumulation in the spinal cord lesion and long-term functional recovery. We performed a moderate thoracic contusion SCI in male and female mice and assessed long-term functional recovery of the hindlimbs using the Basso Mouse Scale (BMS). Spinal cords were collected at 35 dpi and intraspinal neutrophil numbers were quantified at the lesion epicenter. Interestingly, linear regression analysis showed that the total neutrophil numbers within the lesion epicenter of male mice at 35 dpi positively correlated with functional outcomes (R^2^ = 0.38, p = 0.03, Fig. 1l), suggesting that neutrophils may be associated with greater recovery following SCI in male mice. No correlation was observed for female mice. These data suggest a novel and sex-dependent role for neutrophils in mediating recovery following SCI and provide additional evidence for sex differences in neutrophil function after injury.

### Depletion of neutrophils impairs long-term functional recovery in a sex-dependent manner

Given the marked sex differences observed in neutrophil responses to SCI, we next wanted to assess the sex-dependent contribution of neutrophils towards long-term functional recovery. We administered anti-Ly6G depletion antibody (clone 1A8) or IgG control antibody (clone 2A3) immediately following thoracic contusion SCI and assessed long-term functional recovery (Fig. 2a). Contrary to the prevailing dogma that neutrophils are detrimental after SCI, antibody-mediated depletion of neutrophils substantially impaired long-term functional recovery following SCI (Fig. 2b). Remarkably, long-term functional impairment with neutrophil depletion was only observed in male mice (Fig. 2c). While we observed an overall effect for neutrophil depletion on long-term white matter tissue sparing across both sexes, multiple comparisons analyses within each sex were not significant between neutrophil depleted and control mice (Fig. 2d), suggesting that tissue sparing alone may not be responsible for the observed impairment of functional recovery in males.

**Figure 2.**
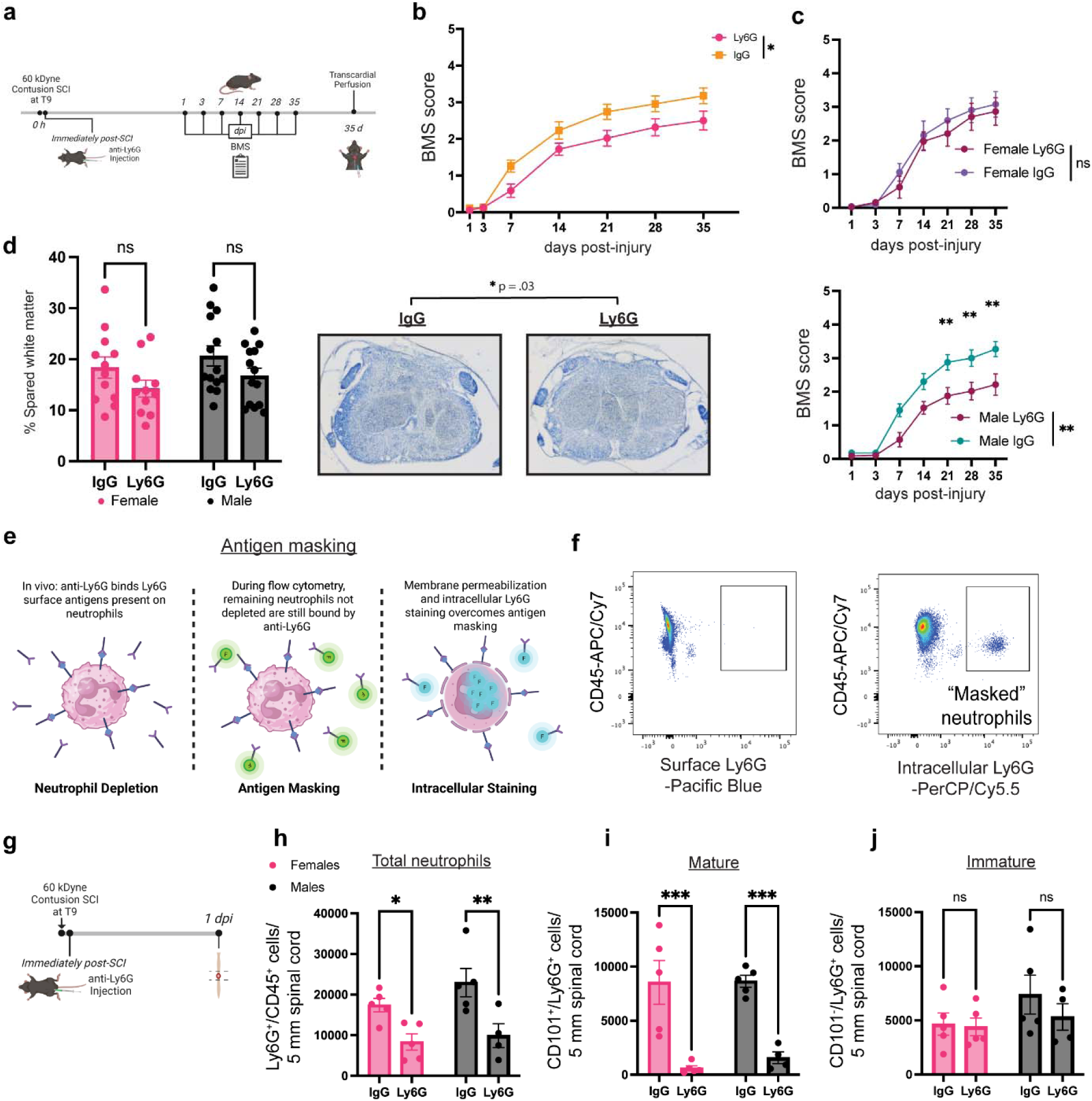
Depletion of mature neutrophils impairs long-term recovery in a sex-dependent manner. (a) Schematic of experimental paradigm for post-SCI depletion utilizing anti-Ly6G antibody (clone 1A8, 2.5 mg/kg) in male and female mice. (b) Depletion of neutrophils immediately after SCI reduced BMS scores. n=13-14/sex/treatment. Repeated measures three-way ANOVA. Main effect of treatment, p = .025. (c) Disaggregation of BMS scores in (b) by sex. Depletion of neutrophils did not alter BMS scores for female mice. Neutrophil depletion reduced BMS scores after SCI in male mice. N=13-14/treatment. Repeated measures two-way ANOVA with Bonferroni’s multiple comparison test. (d) Quantification and representative images of white matter sparing determined by eriochrome cyanine staining. White matter sparing was reduced with neutrophil depletion across both sexes (p = 0.03), however, no difference was observed within each sex. Two-way ANOVA with Sidak’s multiple comparisons test. n=11-14/sex/treatment. (e) Schematic of antigen masking circumvented by intracellular Ly6G staining. (f) Flow cytometry analysis of neutrophil populations with surface (left) and intracellular (right) Ly6G staining after neutrophil depletion. Intracellular Ly6G labels remaining masked neutrophils not labeled with conventional surface staining of Ly6G. (g) Schematic of SCI, depletion of neutrophils, and tissue collection for flow cytometry analysis. (h) Anti-Ly6G antibody administration reduces total intraspinal neutrophils (Ly6G^+^ cells percentage out of total CD45^+^ population) at 1 dpi. (i) Anti-Ly6G antibody administration substantially reduces intraspinal mature (CD101^+^) neutrophil counts. (j) Intraspinal immature (CD101^−^) neutrophils were not altered by anti-Ly6G antibody administration. N = 4-5/sex/treatment. Two-way ANOVA with Sidak’s multiple comparisons test. *p<0.05, **p<0.01, ***p<0.001. Abbreviations: SCI = spinal cord injury, BMS = Basso Mouse Scale, dpi = days post-injury. Mean +/− SEM.

To ensure that anti-Ly6G antibody administration after SCI abrogates intraspinal neutrophil accumulation, we first validated our method for detecting remaining neutrophil populations. Staining for neutrophils that remain following administration of anti-Ly6G antibody can be difficult due to the recently described masking of the Ly6G surface antigen^49^, which can be circumvented by intracellular Ly6G staining (Fig. 2e). In our hands, staining for intracellular Ly6G was effective at labeling remaining neutrophil populations that could not be labeled with conventional surface Ly6G staining techniques following anti-Ly6G administration (Fig. 2f). We performed flow cytometry analysis on spinal cord samples collected 1 day after SCI and neutrophil depletion (Fig. 2g). Anti-Ly6G depletion antibody administered immediately after SCI reduced total intraspinal neutrophil numbers at 1 dpi independent of sex (Fig. 2h). Interestingly, neutrophil depletion substantially abrogated the accumulation of mature (CD101^+^) neutrophils at 1 dpi in both sexes but did not alter immature (CD101^−^) neutrophil accumulation (Fig 2i-j). No differences in the mean fluorescent intensity (MFI) of surface Ly6G were observed between immature and mature circulating neutrophils in uninjured mice (Fig. S2a), suggesting that surface Ly6G abundance alone does not account for in the greater depletion efficacy of mature versus immature neutrophils in the acutely injured spinal cord. Our data suggest that the sex-dependent effects of neutrophil depletion on long-term functional recovery may be due to inherent sex differences in the role of neutrophils in the injured spinal cord rather than differences in the efficacy of neutrophil depletion.

### Intraspinal neutrophil phenotype and putative function are highly dynamic after SCI

To better understand how intraspinal neutrophil phenotype and function may contribute to the sex-dependent functional impairments observed following neutrophil depletion, we analyzed the transcriptional profile of neutrophils isolated from three published scRNAseq datasets^41–43^. Clustering of neutrophils by time revealed a population of intraspinal neutrophils at 3 dpi that appeared distinct from other timepoints (Fig. 3a). For simplicity, neutrophils from sham and uninjured (UN) mice were combined into a single group. A correlation matrix of the transcriptional profiles of neutrophils at different timepoints after SCI confirmed that intraspinal neutrophils at 3 dpi are transcriptionally distinct (Fig. 3b). Analysis of gene expression for neutrophil granule production and metabolism showed drastic differences in transcript levels associated with azurophilic granules, specific granules, and glycolysis across time, further demonstrating that neutrophil phenotype is highly dynamic after SCI (Fig. 3c). In addition, analysis of transcript levels for genes associated with phagocytosis, chemotaxis, and NADPH oxidase (critical for ROS production) shows evidence that neutrophil function is likely highly dynamic following injury.

**Figure 3.**
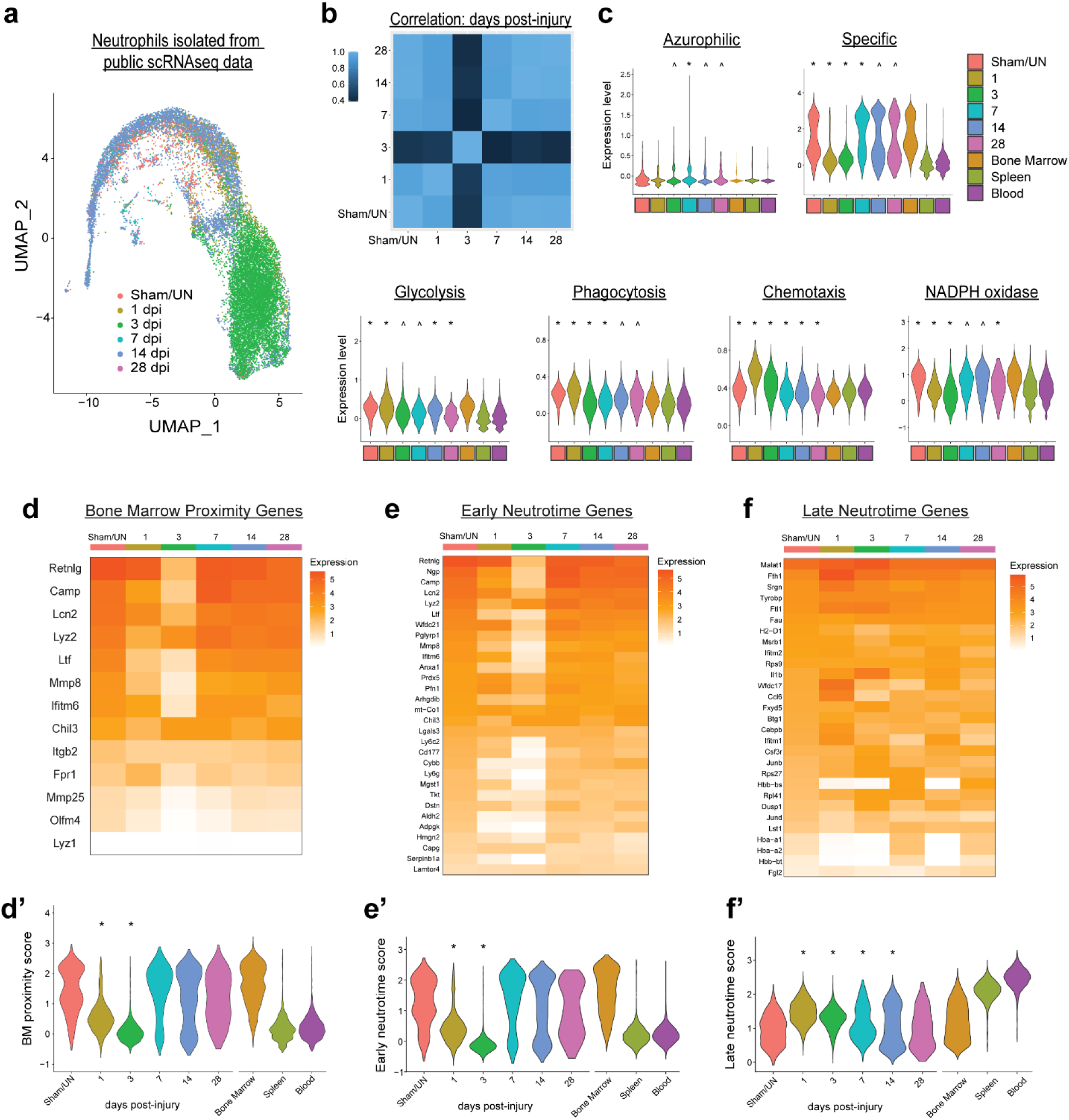
ScRNAseq analysis reveals unique time-dependent neutrophil transcriptional signatures in the injured spinal cord. (a) UMAP of intraspinal neutrophils isolated from three published scRNAseq datasets with thoracic contusion SCI in mice. Neutrophils from sham and uninjured mice are combined together in “sham/UN” group. Neutrophils are transcriptionally distinct across time after SCI. (b) Correlation matrix of the transcriptional profiles of neutrophils at different timepoints after SCI. Neutrophils at 3 dpi are transcriptionally distinct versus all other timepoints. (c) Expression levels of genes associated with azurophilic granules, specific granules, glycolysis, phagocytosis, chemotaxis, and NADPH oxidase (ROS production). (d) Heatmap for genes associated with the bone marrow proximity score. (d’) Quantification of the bone marrow proximity score for neutrophils after SCI. (e) Heatmap for genes associated with the early Neutrotime score. (e’) Quantification of the early Neutrotime score for neutrophils after SCI. (f) Heatmap for genes associated with the late Neutrotime score. (f’) Quantification of the late Neutrotime score for neutrophils after SCI. Neutrophils at 1 and 3dpi express genes indicative of mature neutrophils. Kruskal-Wallis test with Dunn’s Test for post-hoc analysis. *p<0.05 vs. all other timepoints. ^p<0.05 vs. sham/UN. Bone marrow, blood, and spleen neutrophils are not included in analysis. Abbreviations: SCI = spinal cord injury, dpi = days post-injury, BM = bone marrow, ROS = reactive oxygen species.

To determine how neutrophil maturation changes after SCI in the scRNAseq data, we assessed the transcriptional similarity of intraspinal neutrophils to bone marrow neutrophils using the previously described bone marrow proximity score^48^. Intraspinal neutrophils during the acute phase of SCI (1 and 3 dpi) had significantly lower bone marrow proximity scores relative to neutrophils from sham/uninjured animals and all other timepoints post-SCI, suggesting that neutrophils at these timepoints are transcriptionally distinct from the bone marrow neutrophil population (Fig. 3d-d’). By assessing the early and late Neutrotime^44^ scores of intraspinal neutrophils, which quantifies the transcript levels of early and late genes expressed during neutrophil development and maturation (Figs. 3e-e’ and 3f-f’), we confirmed that intraspinal neutrophils at 1 and 3 dpi had uniquely greater maturation profiles when compared to intraspinal neutrophils at other timepoints after SCI, as well as neutrophils from sham and uninjured mice. While intraspinal neutrophils at 1 and 3 dpi both had relatively high mature neutrophil scores, neutrophils at 3 dpi had remarkably unique transcriptional signatures, suggesting functionally distinct properties for these cells.

### Mature neutrophils adopt a pro-resolving phenotype and limit macrophage accumulation in a sex-dependent manner

Since anti-Ly6G antibody administration mitigated the accumulation of mature neutrophils in the acutely injured spinal cord, we performed a gene ontology analysis of differentially expressed genes from neutrophils at 3 dpi relative to neutrophils at all other timepoints. Efferocytosis (mmu04148) was identified as a prominent pathway for neutrophils at 3 dpi, potentially indicating the production of “eat me” signals from neutrophils undergoing phagocytosis (Fig. 4a). It has been previously shown that macrophage phagocytosis of apoptotic neutrophils can shift macrophages towards an anti-inflammatory phenotype and induce a pro-resolution cascade that can promote wound healing and tissue repair^9,10,50^. To further analyze the potential role of neutrophils in resolution of inflammation, we focused on the data from Wang and colleagues^43^, where scRNAseq was performed on leukocytes (CD45^+^ cells) sorted, enriched, and pooled from the injured spinal cords of male and female mice. Analysis of the ligand-receptor interaction strength of neutrophils at 3 dpi revealed high incoming and outgoing interactions between neutrophils and macrophages, as well as monocytes, within the injured spinal cord (Fig. S3a-b). Interaction of neutrophils with macrophages waned by 14 dpi, suggesting that the 3 dpi neutrophil population might play a unique role in regulating macrophage responses to SCI.

**Figure 4.**
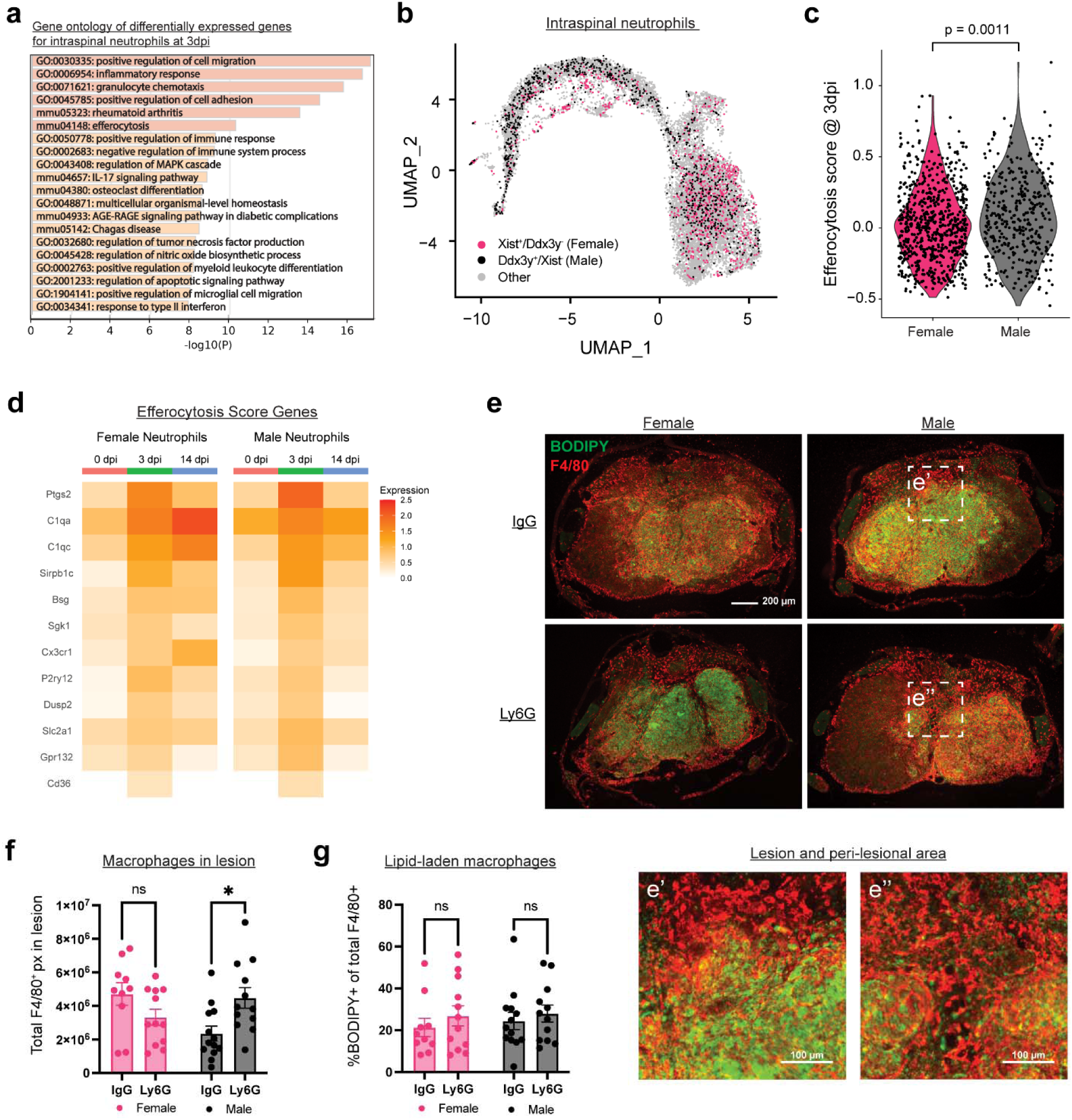
Neutrophils mediate resolution of inflammation after SCI in a sex-dependent manner. (a) Gene ontology (GO) analysis of differentially expressed genes in intraspinal neutrophils at 3 dpi vs. all other timepoints after SCI. (b) UMAP of neutrophils isolated from Wang *et al*. 2022. Female and male neutrophils were identified by expression of transcripts for *Xist* and *Ddx3y* respectively. (c) Quantification of transcript levels for genes associated with the efferocytosis GO term pathway for female and male neutrophils at 3 dpi. (d) Heatmap of gene expression for efferocytosis GO term pathway for neutrophils at 0, 3, 14 dpi SCI. (e) Representative images showing BODIPY labeling of neutral lipids (green) and immunostaining of F4/80 for macrophages (red) in the spinal cord lesion epicenter at 35 dpi in female and male mice with neutrophil depletion (Ly6G) or control (IgG) antibody. e’ and e” are magnified insets. (f) Quantification of the total F4/80^+^ pixels accumulated from seven serial sections spanning 1.5 mm centered on the lesion. Neutrophil depletion exacerbates long-term macrophage accumulation in male, but not female mice. n=10-13/sex/treatment. (g) Quantification of the total BODIPY^+^ pixels out of total F4/80^+^ pixels from seven serial sections spanning 1.5 mm centered on the lesion. No differences in lipid-laden macrophages were observed. Two-way ANOVA with Sidak’s multiple comparison test. *p<0.05. Abbreviations: SCI = spinal cord injury, dpi = days post-injury. Mean ± SEM.

To assess sex differences in the neutrophil transcriptional profiles, we identified male and female neutrophils from the pooled samples by the expression of *Xist* (female) and *Ddx3y* (male) as previously described^38^ (Fig. 4b). We observed greater transcript levels for genes associated with efferocytosis in male neutrophils relative to females (Fig. 4c-d). Since neutrophil efferocytosis can influence subsequent macrophage responses, we assessed the effect of neutrophil depletion on macrophage accumulation at 35 dpi. Neutrophils have also been shown to play a role in regulating the formation and function of myelin lipid-laden (“foamy”) macrophages, therefore, we also assessed lipid-laden macrophage formation and accumulation in the injured spinal cord. While we did not observe any effect of neutrophil depletion on total lipid droplet abundance or lipid-laden macrophage accumulation, we did observe a striking increase in total macrophage accumulation in the injured spinal cords of male, but not female, mice at 35 dpi (Fig. 4e-g). These findings provide evidence for a novel and sex-dependent role of mature neutrophils in promoting resolution of inflammation in the injured spinal cord.

Next, we sought to characterize the effect of neutrophil depletion on other anatomical hallmarks of SCI such as vascular damage, reactive gliosis, and axon sprouting. Neutrophil depletion immediately after SCI did not appear to alter acute blood-spinal cord barrier disruption at 1 dpi (Fig. S4a-b) or the abundance of intraspinal vasculature at 7 or 35 dpi (Fig. S5a-d). We also observed no effects of neutrophil depletion on GFAP expression in the injured spinal cord (Fig. S6a-b). Lastly, as macrophages have recently been implicated in serotoninergic (5-HT^+^) axon sprouting into the lesion following SCI^51^, we assessed the effect of neutrophil depletion on 5-HT^+^ axons at 35 dpi. No differences in 5-HT axon density were observed in the injured spinal cord in either sex (Fig. 5a-b). While acute neutrophil depletion did not impair the accumulation of later arriving neutrophils (Fig. 5c), the relationship between chronic intraspinal neutrophil accumulation and 5-HT^+^ axons was remarkably altered in a sex-dependent manner (Fig. 5d-e). For female mice that received the anti-Ly6G depletion antibody and SCI, neutrophil accumulation at 35 dpi positively correlated with 5-HT axon density (R^2^ = 0.54, p = .01, Fig. 5d) whereas for neutrophil depleted male mice, we observed a trending negative correlation (R^2^ = 0.23, p = .09, Fig. 5e). These data further indicate that acute neutrophil depletion alters the chronic intraspinal inflammatory environment in a sex-dependent manner.

**Figure 5.**
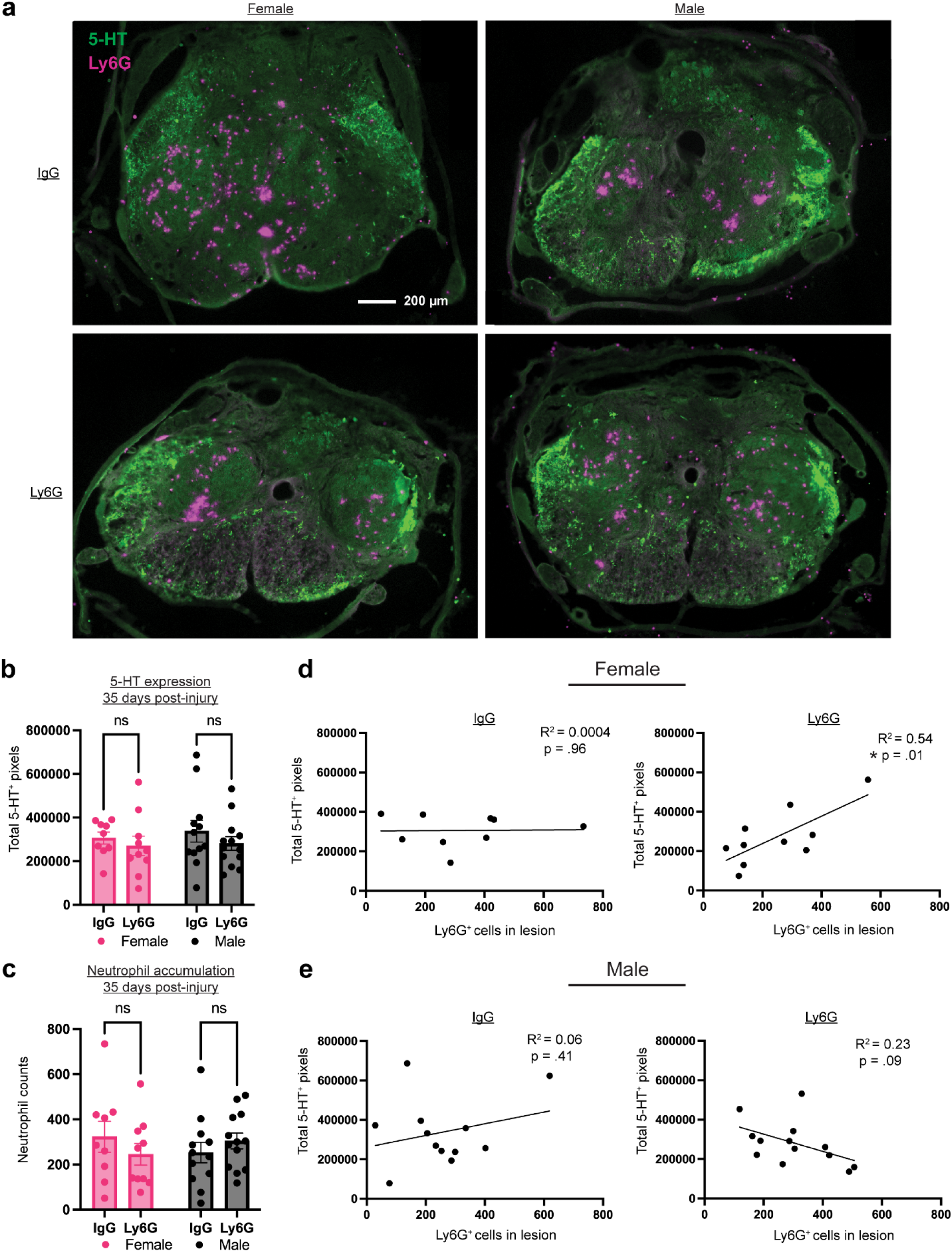
Neutrophil depletion alters the correlation between chronic neutrophil accumulation and 5-HT axon density in a sex-dependent manner. (a) Representative images of 5-HT^+^ axons (green) and accumulated neutrophils (purple) in the lesion epicenter at 35 dpi in male and female mice with neutrophil depletion (Ly6G) or control (IgG) antibody. (b) Quantification of total 5-HT^+^ pixels accumulated from 5 serial sections spanning 1mm centered on the lesion. No differences in 5-HT axon density were observed. n=9-13/sex/treatment. (c) Quantification of total neutrophils (Ly6G^+^ cells) from 5 serial sections spanning 1mm centered on the lesion. No differences in neutrophil accumulation were observed at 35 dpi. n=9/13/sex/treatment. Two-way ANOVA with Sidak’s multiple comparisons test. Mean ± SEM. (d-e) Correlation of total 5-HT^+^ pixels and total neutrophils (Ly6G^+^ cells) from 5 serial sections spanning 1mm centered on the lesion in female (d) and male (e) mice with neutrophil depletion (Ly6G) or control (IgG) antibody. Neutrophil abundance positively correlated with 5-HT axon density in females and trended towards a negative correlation in males. Linear regression analysis. n=9-13/sex/treatment. *p<0.05. Mean ± SEM.

### Pre-injury depletion of neutrophils does not alter functional recovery in either sex

In past studies, neutrophil depletion has been commonly initiated up to 24 hours prior to SCI. To determine if the sex-dependent effects of neutrophil depletion were also time-dependent, we administered anti-Ly6G depletion antibody or IgG control antibody in adult male and female mice at 1 day prior to SCI (Fig 6a). Surprisingly, no significant differences in long-term functional recovery were observed in either sex with pre-injury neutrophil depletion (Fig 6b). Consistent with this finding, pre-injury neutrophil depletion also did not alter long-term white matter sparing (Fig. 6c-d).

**Figure 6.**
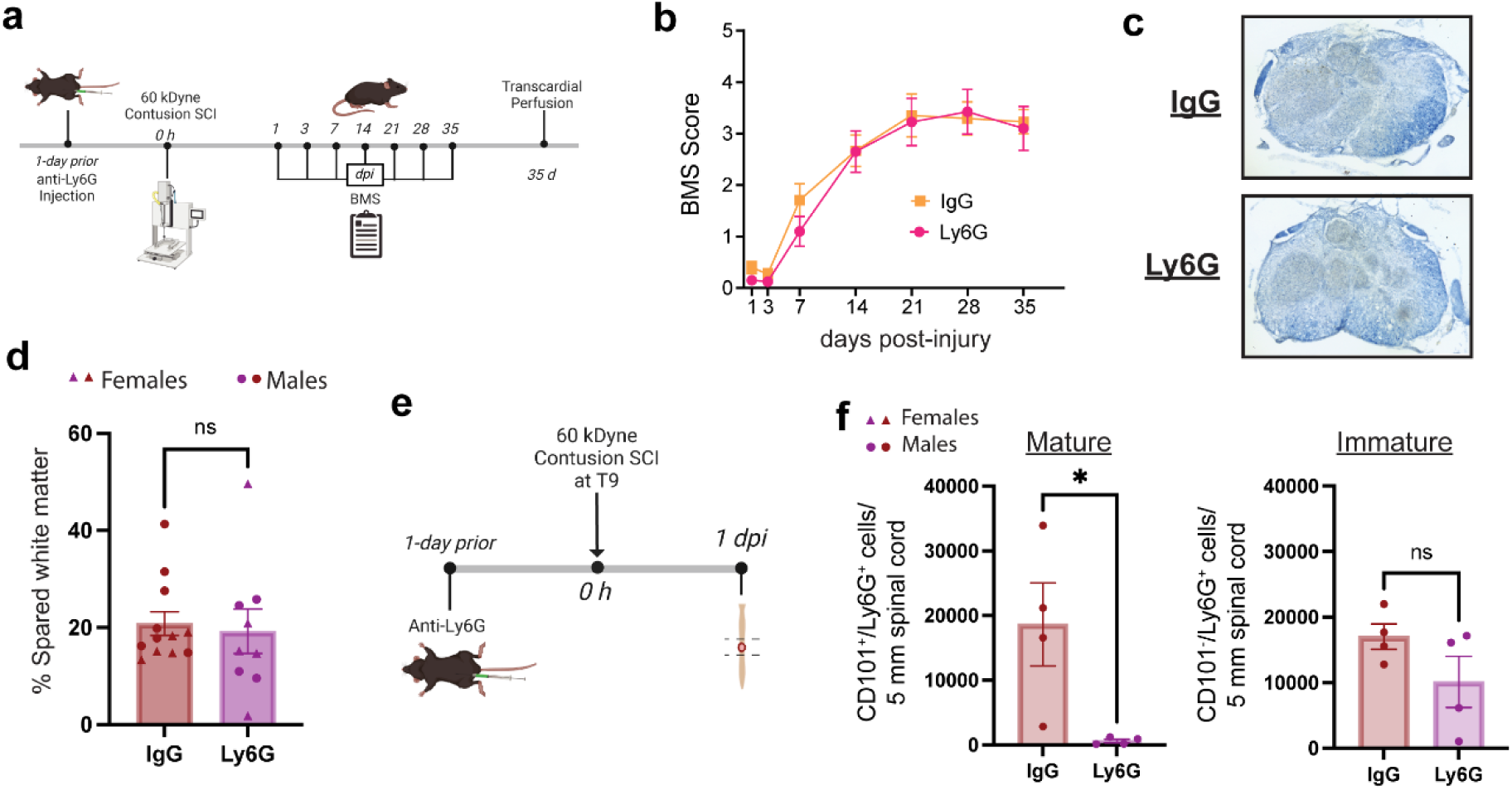
Depletion of mature neutrophils prior to SCI does not impair long-term recovery. (a) Schematic of experimental paradigm for pre-SCI neutrophil depletion. (b) Depletion of neutrophils at 1 day prior to SCI does not alter long-term BMS scores. n=5-7/sex/treatment. Repeated measures three-way ANOVA. (c) Representative images of the lesion epicenter stained with eriochrome cyanine for pre-injury neutrophil depletion (Ly6G) and control (IgG) animals. (d) Quantification of spared white matter at the lesion epicenter for male and female mice at 35 dpi. No sex differences or effect of pre-injury neutrophil depletion were observed on long-term white matter sparing. n=4-7/sex/treatment. Unpaired two-tailed Student’s t test. (e) Schematic of pre-injury depletion of neutrophils, SCI, and tissue collection for flow cytometry analysis. (f) Anti-Ly6G antibody administration substantially reduces intraspinal mature (CD101^+^) neutrophils, but not intraspinal immature (CD101^−^) neutrophils. n=4/treatment. Unpaired two-tailed Student’s t test. Abbreviations: SCI = spinal cord injury, BMS = Basso Mouse Scale, dpi = days post-injury. *p<0.05. Mean ± SEM.

We next wanted to ensure that earlier administration of the depletion antibody was still effective at reducing intraspinal neutrophil accumulation following SCI. Antibody-mediated depletion of neutrophils starting 1-day prior to injury successfully reduced the accumulation of mature neutrophils in the acutely injured spinal cord (Fig. 6e-f). Consistent with post-SCI depletion, pre-injury administration of the anti-Ly6G depletion antibody did not alter the accumulation of immature neutrophils. Collectively, our findings demonstrate that the sex-dependent role of mature neutrophils in mediating long-term recovery is also time-dependent.

## Discussion

Neutrophils have historically been thought to exacerbate secondary tissue damage and worsen long-term outcomes following SCI. Neutrophils respond to injury and infection via cytotoxic effector functions essential for mitigating microbial spread, but often with bystander damage to nearby healthy tissue^4,5,8,52,53^. Limiting neutrophil accumulation in the acutely injured spinal cord, by blocking adhesion receptors or chemoattractants, has commonly led to tissue sparing and greater functional recovery^21,54,55^. However, neutrophil depletion in the context of SCI has produced conflicting results^20–24^ and there is an emerging body of literature demonstrating beneficial roles for neutrophils in CNS and peripheral nerve injury. The newly discovered reparative properties of neutrophils, as well as sex differences in neutrophil function, warrant re-examining the sex-dependent role of neutrophils in the injured spinal cord.

The contribution of neutrophils in SCI is likely multifaceted, with neutrophils having both harmful and reparative functions. Neutrophil heterogeneity may mediate this functional divergence, which could be further complicated by sex differences in neutrophil function. Here we show that neutrophils in the periphery and lesion site rapidly change in number and phenotype after SCI in a time- and sex-dependent manner. We demonstrate that neutrophil depletion worsens long-term functional recovery in male mice, but not females. We demonstrate that neutrophils acquire a mature and inflammation-resolving phenotype in the acutely injured spinal cord and that depletion of mature neutrophils exacerbates long-term inflammation in a sex-dependent manner. While neutrophil depletion does not appear to alter other classic anatomical hallmarks of SCI, it does shift the long-term relationship between later neutrophil accumulation and 5-HT^+^ axons density in a sex-dependent manner. Our findings demonstrate that neutrophils respond to SCI in a time- and sex-dependent manner and engage in sex-specific resolution of inflammation and potentially other beneficial activities that are critical for mediating long-term functional recovery.

During maturation, neutrophils will increase CXCR2 expression and downregulate CXCR4 to facilitate release from the bone marrow into circulation^30^. Neutrophil mobilization after injury is coordinated in part by an increase in granulocyte colony-stimulating factor (G-CSF), which stimulates the downregulation of CXCR4 to enhance neutrophil release into circulation^56^. While sex differences have been previously shown for circulating and bone marrow neutrophils^27,36,38,39^, our study is among the first to demonstrate sex differences in neutrophil mobilization as well as granulopoiesis after injury. Neutrophil abundance after SCI has been shown to be predictive of long-term functional recovery^57^, which may be reflective of injury-induced neutrophilia.

To the best of our knowledge, our study is the first to demonstrate that the maturation profile of intraspinal neutrophils changes over time in a sex-dependent manner. Interestingly, the time course of the maturation profile for intraspinal neutrophils is disparate from that of the circulating population. It is possible that, in addition to infiltrating from circulation, neutrophils may infiltrate into the injured spinal cord along other routes contingent on maturation status. A recent study found that skull and vertebral bone marrow neutrophil populations were capable of bypassing circulation and rapidly respond to inflamed CNS tissue by migrating through microscopic channels connecting the bone marrow lumen with the meninges^58^. This finding is concordant with our flow cytometry data and scRNAseq analyses showing that, in uninjured or sham mice, immature neutrophils still associate with the spinal cord, and we see this at later time points after 7 dpi as well. By migrating directly from the bone marrow, immature neutrophils may be able to enter the spinal cord meninges and possibly the tissue parenchyma, thus allowing for the disconnect between circulating and intraspinal neutrophil maturation states. The mechanisms driving the association of immature neutrophils with the uninjured and chronically-injured spinal cord remain unclear and should be addressed in future studies.

In rodent models of SCI, neutrophil accumulation has been shown to peak within 1 dpi and substantially declines soon after^20,21,24^. We show that neutrophil accumulation in the injured spinal cord is greater in male mice relative to females, which may be due to the increased neutrophil mobilization into the blood after injury. By pulsing actively dividing neutrophil progenitors with EdU prior to injury, we demonstrate that neutrophils survive in the SCI lesion site for at least 5 weeks after injury, whereas neutrophils in the blood and bone marrow appear to retain their normal short lifespan. While extended neutrophil lifespan is generally accepted in normal tissue environments and has been recently demonstrated in tumors^59,60^, we believe our study is the first to demonstrate prolonged neutrophil survival (several weeks) in a tissue injury site. Furthermore, we demonstrate that newly generated neutrophils pulsed with EdU several weeks after SCI can still enter the SCI lesion site despite the marked reduction in neutrophil numbers. To the best of our knowledge, this is the first study to identify late-arriving neutrophils in a rodent model of SCI.

Neutrophil depletion in thoracic contusion models of SCI has produced varied effects on long-term outcomes ranging from impaired recovery^20^, to no effects^22,23^, to marked neuroprotection^21^. Early studies utilized the GR-1 depletion antibody, which can target cells expressing Ly6G and Ly6C, including monocytes^24^. However, it is important to note that only a single sex was considered in these studies. In fact, the only study to use male mice (sex was not published, but was confirmed by Dr. David Stirling) saw worse long-term functional recovery with neutrophil depletion, albeit with the GR-1 antibody^20^. The follow-up studies utilized female mice and mostly found minimal effects on long-term recovery, particularly when using the Ly6G depletion antibody (clone 1A8). While many methodological differences exist between these studies, our study suggests that the inconsistent effects can be partially explained by sex- and time-dependent effects of neutrophil depletion after SCI.

Reparative roles for neutrophil subsets in models of sterile injury have largely been attributed to immature neutrophils. Within the CNS, immature neutrophils have been shown to promote neuron survival, promote axon regeneration in the injured spinal cord, and contribute to regenerative and neuroprotective processes^28^. In our study, we provide novel evidence of a sex-dependent reparative role for mature neutrophils in CNS injury. Neutrophils are capable of synthesizing many specialized pro-resolving mediators that work to regulate leukocyte accumulation and phenotype. Neutrophil extracellular traps can sequester cytokines and chemokines and matrix metalloproteinase-9, released from neutrophil granules, can degrade damage associated molecular patterns to mitigate further leukocyte recruitment^9,61,62^. In addition, apoptotic neutrophils can induce resolution of inflammation by promoting an anti-inflammatory phenotype in phagocytosing macrophages during efferocytosis^9,10,50,63^. Our data indicate a direct interaction of neutrophils with macrophages, including the expression of genes involved in efferocytosis, which likely collectively work with indirect mechanisms (sequestration and degradation of immune stimulating cues) to mitigate subsequent inflammation. We focused on macrophages due to their prevalence in the SCI lesion site, however, future work should examine the effect of neutrophils on other leukocyte subsets, including lymphocytes. Due to recent evidence that intraspinal macrophages negatively regulate axonal sprouting^51^, we anticipated reduced 5-HT^+^ axons in and near the lesion site in male mice following neutrophil depletion and the associated increased accumulation of macrophages. While neutrophil depletion did not appear to alter axonal sprouting, there was a sex-dependent change in the correlation of neutrophil numbers and 5-HT^+^ axonal density in the chronically injured spinal cord. These data suggest that long-term neutrophil phenotype may be altered by acute depletion with beneficial and detrimental effects in female and male mice, respectively.

While other studies on neutrophil heterogeneity have found mature and immature neutrophil subsets to have differing levels of Ly6G expression^25,28,33^, we found that circulating mature and immature neutrophils equally express the Ly6G surface antigen. Additionally, we demonstrate that anti-Ly6G administration in the context of SCI specifically abrogates mature neutrophil accumulation in the injured spinal cord in both male and female mice. A similar effect has been previously reported on splenic neutrophils with anti-Ly6G administration, where again the mature neutrophil population was predominantly depleted by anti-Ly6G^33^. Additionally, because we only administer one acute dose of anti-Ly6G in both of our depletion studies, the depletion capacity of anti-Ly6G in our experiments is inherently transient. Therefore, the pre-injury anti-Ly6G antibody administration might not deplete neutrophils long enough to significantly impact the later intraspinal population of mature neutrophils (e.g. 3 dpi) to the same extent as post-injury anti-Ly6G administration. Interestingly, one study that employed the anti-Ly6G antibody for neutrophil depletion in the context of SCI previously showed that a single dose of anti-Ly6G was effective at depleting neutrophils for 3 days in uninjured female mice^22^. While we observe no effect of acute neutrophil depletion on later neutrophil accumulation at 35 dpi, the duration of neutrophil depletion after SCI in both sexes remains to be determined.

Collectively, our results identify sex differences in neutrophil responses to SCI and demonstrate that mature neutrophils in the acutely injured spinal cord are predominantly beneficial for long-term functional recovery in male mice. To the best of our knowledge, our study is the first to demonstrate sex-dependent roles for mature neutrophils in resolution of inflammation after injury. Our work also addresses many of the inconsistent effects previously observed in past neutrophil depletion studies in SCI by demonstrating the sex- and time-dependent effects of neutrophils in long-term functional recovery. Future studies are needed to determine why resolution of inflammation by neutrophils only occurs in male mice after SCI and if these processes can be augmented for future sex-specific strategies to promote long-term recovery after SCI.

## Methods

### Mice

All procedures were performed in accordance with protocols approved by the Institutional Animal Care and Use Committee at Texas A&M University. All experiments used adult male and female C57Bl/6J (WT) mice (age 12-20 weeks; age-matched within experiments) ordered from The Jackson Laboratory and maintained in our colony. Mice were housed in groups of two to five prior to antibody administration and surgery. Following antibody administration or injury, female mice were housed in groups of two to three, and male mice were single-housed. Mice were housed in a climate-controlled room on a 12 h light/dark cycle with food and water provided ad libitum.

### Antibody-mediated neutrophil depletion

Animals were randomly assigned to groups and experimenters were blinded to treatment throughout the duration of each experiment. Animals received either anti-Ly6G (2.5mg/kg, clone 1A8, InVivoPlus, BioXCell) or the control non-specific IgG antibody (2.5mg/kg, clone 2A3, InVivoPlus, BioXCell) via intraperitoneal (i.p.) injection. Antibodies were diluted in sterile InVivoPure dilution buffer (pH=7.0, BioXCell) and prepared in a sterile bio-safety cabinet. For studies assessing the effect of neutrophil depletion on the recovery process following SCI, mice were injected either 1-day prior to injury or immediately post-injury following closing of the skin incision.

### Spinal cord injury

A moderate spinal cord contusion injury was performed for all studies. Mice were anesthetized using 2% isoflurane, a laminectomy was performed at thoracic vertebral level nine (T9), and a 60-kilodyne (kDyne) contusive SCI with a one-second dwell time was administered using the Infinite Horizon Impactor. Following the SCI, 50 µl of 0.25% bupivacaine was administered directly onto the sutured musculature post-injury prior to closing the skin incision with wound clips, then animals received 1 mL of saline via subcutaneous (s.q.) injection. Animals were kept on a circulating water pad set at 37°C to maintain body temperature before and after surgery.

### Post-operative care

To ensure hydration and to prevent urinary tract infections, mice were given saline (1 mL, s.q.) and Baytril (2.5mg/kg) for the first 10 days post-injury. Food pellets were placed on the floor of the cages for easy access as animals recovered. Manual bladder expression was performed twice daily until euthanasia. Body weight was measured at 1 and 3 days post-injury (dpi) and then weekly until euthanasia. Animals with significant reductions in body weight (>15%) were given Nutrical on food pellets to encourage eating.

### Open field locomotor testing

The Basso Mouse Scale (BMS) was performed at 1, 3, 7, 14, 21, 28, and 35 days post-SCI to assess hindlimb locomotor recovery^40^. The 9-point BMS test assesses multiple aspects of hindlimb function such as the degree of hindlimb movement, paw placement, stepping, and coordination where higher scores indicate greater levels of hindlimb functional recovery. For BMS testing, animals were placed into a rectangular open field and were rated by two independent scorers for 4-minutes. Scorers were blinded to treatment and group throughout the duration of the study.

### Flow cytometry

Mice were anesthetized by i.p. injection of 2.5% avertin (0.02 mL/g body weight) and bilateral thoracotomy was performed. Peripheral blood was collected via cardiac puncture of the right ventricle using an insulin syringe containing 200 μL of 3.6 mg/mL EDTA (Invitrogen) to prevent clotting. Blood was transferred into a microcentrifuge tube, placed on ice, and additional EDTA (3.6 mg/mL) was added till a 1:1 ratio of blood to EDTA was reached. Blood/EDTA solution was then transferred into a 15 mL conical containing 10 mL of ice cold HBSS (without calcium, magnesium, or phenol red). Following collection of peripheral blood, approximately 5 mm of spinal cord tissue (including meninges) centered on the injury site was rapidly dissected and placed into a microcentrifuge tube containing 300 μL RPMI media on ice. While kept on ice, the spinal cord tissue was mechanically dissociated using a plastic tissue pestle. The solution was triturated before being filtered through a 70-μm nylon mesh cell strainer. The filtered solution was added to 10 mL of cold HBSS on ice. Following collection of the spinal cord, the femur and tibia were dissected from one of the hindlimbs. Both bones were then transferred to 5 mL of HBSS on ice. The lateral and medial condyles of the femur were removed as well as the femoral head using small dissecting scissors, and 10 mL of ice cold HBSS was flushed through the lumen to flush the cells into an empty 50mL conical. The sample was then triturated on ice with a 19G needle to evenly suspend the bone marrow cells in HBSS.

Blood, bone marrow, and spinal cord samples were centrifuged at 500 × g, 4°C for 5 min. Pellets were then lysed with 10 mL (blood) or 1 mL (bone marrow & spinal cord) of 1X red blood cell (RBC) lysis buffer (BD Pharm Lyse) at 4°C. Blood was lysed for 6 min while bone marrow and spinal cord were lysed for 5 min. Blood samples were quenched with 40 mL ice cold FACS buffer (1 mM EDTA and 2% fetal bovine serum in HBSS). Bone marrow and spinal cord samples were quenched with 10 mL ice cold FACS buffer. Samples were centrifuged at 400 × g, 4°C for 5 min, pellets were resuspended in 10 mL cold HBSS, and 1×10^6^ bone marrow cells were transferred to a new 15 mL conical. Samples were centrifuged again, supernatant was tipped off, and the pellets were resuspended in Zombie Red Fixable Viability Kit (Biolegend; 1:500) for 30 min at 4°C for live/dead cell staining. Samples were quenched with 10 mL cold FACS buffer and centrifuged. Supernatant was tipped off and samples were resuspended in remaining supernatant and transferred to a 96-well round bottom plate. Samples were centrifuged at 800 × g, 4°C for 2 min, the supernatant was discarded, and the pellets were incubated with anti-mouse CD16/32 Fc blocking antibody (1:100; Biolegend, cat: 101302) at 4°C for 20 min. Samples were centrifuged, supernatant discarded, and pellets were resuspended in 200 μL cold FACS buffer. For isotype staining, 100 μL of cell solution was split into a separate well and samples were centrifuged again.

Pellets were then incubated for 30 min at 4°C with a cocktail of the following antibodies: Pacific Blue-conjugated rat anti-mouse Ly6G (1:200; 1A8, Biolegend, cat: 127612), FITC-conjugated rat anti-mouse/human CD11b (1:200; M1/70, Biolegend, cat: 101206), APC-Cyanine7-conjugated rat anti-mouse CD45 (1:200; 30-F11, Biolegend, cat: 103115), and PE-Cyanine7-conjugated rat anti-mouse CD101 (1:100; Moushi101, Invitrogen, ref: 25-1011-82). PE-Cyanine7-conjugated rat IgG2α,κ (1:100; RTK2758, Biolegend, cat: 400521) served as an isotype control for CD101. Samples were quenched with 200 μL of cold FACS and centrifuged, supernatant discarded, and then resuspended in FACS and centrifuged again. For fixation and permeabilization of the cell membranes for intracellular Ly6G staining, pellets were then incubated with 100 μL of BD Fixation/Permeabilization solution (Biolegend) for 20 min at 4°C. Samples were washed with 200 μL of 1Χ BD Perm/Wash buffer (Biolegend), centrifuged, resuspended in cold FACS buffer, and stored at 4°C overnight. The following day, cells were centrifuged at 800 × g at 4°C for 2 min and pellets were resuspended in 1Χ BD Perm/Wash buffer containing PerCP-Cyanine5.5-conjugated rat anti-mouse Ly6G (1A8, Biolegend, cat: 127616) for intracellular Ly6G staining. For EdU labeling, cells were resuspended in 1X Click-iT perm/wash reagent for 15 minutes in the dark. 50uL of resuspended cells were moved to a separate well for control staining. Cells were centrifuged and resuspended in the reaction cocktail from the Click-iT EdU Alexa Fluor 488 Flow Cytometry Assay Kit (Invitrogen C10632) with 20% of the recommended amount of azide. Control wells received ddH20 instead of azide. Cells were washed and centrifuged twice, then resuspended in FACS buffer.

Flow cytometry was performed on the cell suspensions using a BD Fortessa X-20. For spinal cord samples, 10 μL of Accucheck Counting Beads (Invitrogen, ref: PCB100) were added to the cell suspension before the sample was run through the cytometer. At least 50,000 events for blood and bone marrow and all events for spinal cord samples were collected. Compensation beads were generated using ArC Amine Reactive Bead Compensation Kit (Invitrogen) for Zombie Red and AbC Anti-Rat/Hamster Bead Kit (Invitrogen) for all other staining antibodies. Data analysis was performed using FlowJo software. CD101^+^ populations were gated using the CD101 isotype signal out of Ly6G^+^ cells.

### White matter sparing

Animals were euthanized at 35-days post-SCI and transcardially perfused with 25 mL of ice cold 1X PBS followed by 35 mL of cold 4% paraformaldehyde in PBS. Spinal columns were dissected and post-fixed in 4% paraformaldehyde overnight at 4°C. Spinal cords were then dissected and cryoprotected in 30% sucrose with 0.2% sodium azide. After cryoprotection, 4 mm of spinal cord tissue centered over the injury site was dissected and embedded in Tissue-Tek optimal cutting temperature medium (Sakura) and rapidly frozen on dry ice. Transverse tissue sections (25-μm thick) were cut in series along the rostral-caudal axis and every tenth section was selected for eriochrome cyanine staining. Images were captured using a Leica DM 6B microscope. Total area of residual myelin was quantified using ImageJ and the section with the least white matter area was selected as the lesion epicenter.

### Immunofluorescent and neutral lipid staining

Spinal cords and transverse sections were collected as described for white matter sparing. Slides with transverse tissue sections were dried on a plate warmer set at 37°C for 10 min then washed with 1X PBS at room temperature (RT) for 5 min. Sections were incubated in blocking solution for 1 hour at RT in humidified chambers to prevent drying. For F4/80 and BODIPY co-labeling, blocking solution consisted of 5% normal donkey serum (NDS) and 0.1% bovine serum albumin (BSA) diluted in 1X PBS. For CD31 immunolabeling, blocking solution consisted of 5% NDS, 5% normal goat serum (NGS), 0.1% BSA, and 0.25% Tween20 in 1X PBS. For GFAP immunolabeling, blocking solution consisted of 5% NDS, 0.1% BSA, and 0.25% Tween20 in 1X PBS. For 5-HT and Ly6G co-immunolabeling, blocking solutions consisted of 5% NDS, 0.1% BSA, and 0.2% Triton in 1X PBS. Next, sections were incubated in blocking solution containing rat anti-mouse F4/80 primary antibody (1:100; BM8, Biolegend), hamster anti-mouse CD31 primary antibody (1:200; Millipore), chicken anti-mouse GFAP polyclonal primary antibody (1:200; Encore), 5-HT primary antibody (1:250; ImmunoStar, cat: 20080) and/or Alexa Fluor 647-conjugated rat anti-mouse Ly-6G (1:200; Biolegend, cat: 127610) overnight at 4°C in humidified chambers. The following morning, sections were brought to RT for 30 min, then washed 3X in 1X PBS. Sections were incubated for 2h at RT in blocking solution containing the appropriate Alexa Fluor dye-conjugated secondary antibodies. For BODIPY labeling, sections were washed 3X in 1X PBS, then incubated in 2 μM BODIPY diluted in 1X PBS for 1 hour at RT. Sections were washed 3X in 1X PBS and coverslipped with Prolong Gold Antifade with DAPI. Images were captured using a Nikon Eclipse upright microscope equipped with a 10x objective (0.45 NA) or 20x objective (0.75 NA). Background fluorescence was captured in an unused channel (Texas Red) to be later used for background subtraction during MATLAB analysis.

### Image analysis

Custom MATLAB code was used to quantify F4/80^+^ pixels for macrophages, BODIPY^+^ pixels for lipid droplets, F4/80^+^BODIPY^+^ pixels for lipid-laden macrophages, CD31^+^ pixels for vasculature, GFAP^+^ pixels for glial scar formation, Ly6G^+^ cells for neutrophils, and 5-HT^+^ pixels for serotonergic fiber sprouting. Sections were manually traced by independent and blinded researchers and values were averaged. F4/80^+^BODIPY^+^ pixels were normalized to the total number of F4/80^+^ pixels in the selected region of interest (ROI). CD31^+^ and GFAP^+^ pixels were normalized to total number of pixels in the selected ROI. For F4/80 and BODIPY co-labeling, a total of 1500 μm spanning the lesion was analyzed and sum total of averaged values was calculated. For CD31 staining at 35-days post-injury, 1250 μm of tissue centered on the lesion epicenter was analyzed. For CD31 staining at 7-days post-injury, we analyzed a 2250 μm of the lesion to encompass potential differences in vasculature more rostral and caudal to the lesion epicenter. For GFAP staining, 1000 μm of lesion was analyzed and sum total of averaged values was calculated. For Ly6G and 5-HT staining, 1000 μm of lesion was analyzed and sum total of averaged values was calculated.

### Single cell RNA sequencing analysis

Publicly available single cell RNA sequencing (scRNAseq) data was collected from recent publications that utilized a lower thoracic contusion SCI in WT mice^41–43^, as well as scRNAseq data on neutrophil populations from the blood, spleen, and bone marrow^44^, using the Sequence Read Archive (SRA) toolkit. The prefetch command was used to locally download SRA files, and the FasterQ-dump command to extract paired FastQ files. Bam files were converted to FastQ files using the bamtofastq tool. Files were renamed to follow Illumina naming conventions, compressed using the gzip command, and uploaded to the 10x Genomics Cloud for alignment with the mouse (mm10) 2020-A transcriptome reference. Raw cell matrices were downloaded to ensure all the neutrophils were captured^45^ and cell matrices were imported into R and processed using Seurat version 4.4.0.

Raw cell matrices were read into R and turned into Seurat objects using the Read10X and CreateSeuratObject functions respectively. Cells were filtered if they contained over 15% of mitochondrial associated genes, less than 90 RNA features, or more than 3000 RNA features. Standard Seurat procedures were followed for normalizing, scaling, clustering, and integrating data across time points. Neutrophils were identified and subset out of the three datasets using the SingleR package^46^ and confirmed with established neutrophil markers (including Ly6g, S100a9, and S100a8). Once isolated, neutrophils from each dataset were integrated together and doublets were identified using the scDblFinder package with default parameters. Gene annotation was done using Metascape^47^. Mus musculus was chosen for both input species and analysis species.

Several gene sets were used to track transcriptional markers of maturity, including bone marrow proximity^48^ and Neutrotime^44^. Scores were calculated using the AddModuleScore function in Seurat. The R package CellChat was used to explore ligand-receptor interactions (Jin 2021).

### Statistical analyses

Statistical analyses were performed using GraphPad Prism (GraphPad Software). For immunostaining and eriochrome cyanine analyses, animals with poor tissue section quality were removed prior to unblinding. Flow cytometry, white matter sparing, immunostaining, and immunoblotting data were analyzed using two-way ANOVA with Sidak’s multiple comparison post hoc test with a 95% confidence interval. Prior to disaggregation by sex, flow cytometry and white matter sparing data were analyzed using an unpaired two-tailed Student’s *t* test. BMS scores were first analyzed using a three-way, repeated measures ANOVA. Where appropriate, BMS scores were disaggregated by sex and analyzed using a two-way, repeated measures ANOVA followed by Bonferroni’s multiple comparison test with a 95% confidence interval. Simple linear regression was performed to assess the relationship between neutrophil accumulation and 5-HT^+^ pixels, as well as neutrophil accumulation and BMS scores. Grubbs’ Test (α = .05) performed in GraphPad Prism identified a single outlier for Ly6G^+^ cells at 35 dpi (female, Ly6G) that was subsequently removed from the analysis. The scRNAseq scores were compared using Kruskal-Wallis test with Dunn’s Test for post-hoc analysis. All R^2^ and p values are reported with the corresponding graphs. For all analyses, significance was designated as *p* < 0.05. Data are reported as the mean ± standard error of the mean (SEM).

## Acknowledgements

The authors would like to thank the Texas A&M University Flow Cytometry & Cell Sorting Facility for the use of their equipment. The authors thank Dr. Jennifer Dulin for the use of the Nikon microscope. The diagrams in Figures 1a, 1f, 2a, 2e, 2g, 6a, 6e, S1b, and S1j were created using Biorender.com. Funding support was provided by Mission Connect, a program of TIRR Foundation, the National Science Foundation Graduate Research Fellowship under Grant No. 2139772, and NIH R01NS122961.

## References

1. Ahuja, C. S. et al. Traumatic spinal cord injury. Nat Rev Dis Primers 3, 17018 (2017).

2. Beck, K. D. et al. Quantitative analysis of cellular inflammation after traumatic spinal cord injury: evidence for a multiphasic inflammatory response in the acute to chronic environment. Brain 133, 433–447 (2010).

3. Stirling, D. P. & Yong, V. W. Dynamics of the inflammatory response after murine spinal cord injury revealed by flow cytometry. Journal of Neuroscience Research 86, 1944–1958 (2008).

4. Ley, K. et al. Neutrophils: New insights and open questions. Science Immunology 3, eaat4579 (2018).

5. Neirinckx, V. et al. Neutrophil contribution to spinal cord injury and repair. Journal of Neuroinflammation 11, 150 (2014).

6. McCreedy, D. A. et al. Spleen tyrosine kinase facilitates neutrophil activation and worsens long-term neurologic deficits after spinal cord injury. J Neuroinflammation 18, 302 (2021).

7. Zivkovic, S., Ayazi, M., Hammel, G. & Ren, Y. For Better or for Worse: A Look Into Neutrophils in Traumatic Spinal Cord Injury. Front Cell Neurosci 15, 648076 (2021).

8. Burn, G. L., Foti, A., Marsman, G., Patel, D. F. & Zychlinsky, A. The Neutrophil. Immunity 54, 1377–1391 (2021).

9. Peiseler, M. & Kubes, P. More friend than foe: the emerging role of neutrophils in tissue repair. Journal of Clinical Investigation 129, 2629–2639 (2019).

10. Wang, J. Neutrophils in tissue injury and repair. Cell Tissue Res 371, 531–539 (2018).

11. Blázquez-Prieto, J. et al. Impaired lung repair during neutropenia can be reverted by matrix metalloproteinase-9. Thorax 73, 321–330 (2018).

12. Harty, M. W. et al. Neutrophil Depletion Blocks Early Collagen Degradation in Repairing Cholestatic Rat Livers. The American Journal of Pathology 176, 1271–1281 (2010).

13. Nishio, N., Okawa, Y., Sakurai, H. & Isobe, K. Neutrophil depletion delays wound repair in aged mice. Age (Dordr) 30, 11–19 (2008).

14. Saijou, E. et al. Neutrophils alleviate fibrosis in the CCl4-induced mouse chronic liver injury model. Hepatol Commun 2, 703–717 (2018).

15. Lindborg, J. A., Mack, M. & Zigmond, R. E. Neutrophils Are Critical for Myelin Removal in a Peripheral Nerve Injury Model of Wallerian Degeneration. J Neurosci 37, 10258–10277 (2017).

16. Horckmans, M. et al. Neutrophils orchestrate post-myocardial infarction healing by polarizing macrophages towards a reparative phenotype. Eur Heart J 38, 187–197 (2016).

17. Gong, Y. & Koh, D.-R. Neutrophils promote inflammatory angiogenesis via release of preformed VEGF in an in vivo corneal model. Cell Tissue Res 339, 437–448 (2010).

18. Alvarenga, D. M. et al. Paradoxical Role of Matrix Metalloproteinases in Liver Injury and Regeneration after Sterile Acute Hepatic Failure. Cells 7, 247 (2018).

19. Kovtun, A. et al. The crucial role of neutrophil granulocytes in bone fracture healing. Eur Cell Mater 32, 152–162 (2016).

20. Stirling, D. P., Liu, S., Kubes, P. & Yong, V. W. Depletion of Ly6G/Gr-1 Leukocytes after Spinal Cord Injury in Mice Alters Wound Healing and Worsens Neurological Outcome. Journal of Neuroscience 29, 753–764 (2009).

21. Brennan, F. H. et al. Complement receptor C3aR1 controls neutrophil mobilization following spinal cord injury through physiological antagonism of CXCR2. JCI Insight 4, e98254.

22. Lee, S. M., Rosen, S., Weinstein, P., van Rooijen, N. & Noble-Haeusslein, L. J. Prevention of Both Neutrophil and Monocyte Recruitment Promotes Recovery after Spinal Cord Injury. Journal of Neurotrauma 28, 1893–1907 (2011).

23. Nguyen, H. X. et al. Systemic Neutrophil Depletion Modulates the Migration and Fate of Transplanted Human Neural Stem Cells to Rescue Functional Repair. J. Neurosci. 37, 9269–9287 (2017).

24. Saiwai, H. et al. Ly6C+Ly6G− Myeloid-derived suppressor cells play a critical role in the resolution of acute inflammation and the subsequent tissue repair process after spinal cord injury. Journal of Neurochemistry 125, 74–88 (2013).

25. Evrard, M. et al. Developmental Analysis of Bone Marrow Neutrophils Reveals Populations Specialized in Expansion, Trafficking, and Effector Functions. Immunity 48, 364–379.e8 (2018).

26. Mackey, J. B. G., Coffelt, S. B. & Carlin, L. M. Neutrophil Maturity in Cancer. Frontiers in Immunology 10, 1912 (2019).

27. Gupta, S. et al. Sex differences in neutrophil biology modulate response to type I interferons and immunometabolism. Proceedings of the National Academy of Sciences 117, 16481–16491 (2020).

28. Sas, A. R. et al. A new neutrophil subset promotes CNS neuron survival and axon regeneration. Nat Immunol 21, 1496–1505 (2020).

29. Aroca-Crevillén, A., Vicanolo, T., Ovadia, S. & Hidalgo, A. Neutrophils in Physiology and Pathology. Annual Review of Pathology: Mechanisms of Disease 19, 227–259 (2024).

30. Adrover, J. M. et al. A Neutrophil Timer Coordinates Immune Defense and Vascular Protection. Immunity 50, 390–402.e10 (2019).

31. Coffelt, S. B., Wellenstein, M. D. & de Visser, K. E. Neutrophils in cancer: neutral no more. Nat Rev Cancer 16, 431–446 (2016).

32. Marini, O. et al. Mature CD10+ and immature CD10-neutrophils present in G-CSF-treated donors display opposite effects on T cells. Blood 129, 1343–1356 (2017).

33. Deniset, J. F., Surewaard, B. G., Lee, W.-Y. & Kubes, P. Splenic Ly6Ghigh mature and Ly6Gint immature neutrophils contribute to eradication of S. pneumoniae. J Exp Med 214, 1333–1350 (2017).

34. Consalvo, K. M., Kirolos, S. A., Sestak, C. E. & Gomer, R. H. Sex-Based Differences in Human Neutrophil Chemorepulsion. The Journal of Immunology 209, 354–367 (2022).

35. Ratajczak-Wrona, W. et al. Sex-dependent dysregulation of human neutrophil responses by bisphenol A. Environmental Health 20, 5 (2021).

36. Blazkova, J. et al. Multicenter Systems Analysis of Human Blood Reveals Immature Neutrophils in Males and During Pregnancy. The Journal of Immunology 198, 2479–2488 (2017).

37. Molloy, E. J. et al. Sex-specific alterations in neutrophil apoptosis: the role of estradiol and progesterone. Blood 102, 2653–2659 (2003).

38. Kim, M., Lu, R. J. & Benayoun, B. A. Single-cell RNA-seq of primary bone marrow neutrophils from female and male adult mice. Sci Data 9, 442 (2022).

39. Lu, R. J. et al. Multi-omic profiling of primary mouse neutrophils predicts a pattern of sex- and age-related functional regulation. Nat Aging 1, 715–733 (2021).

40. Basso, D. M. et al. Basso Mouse Scale for Locomotion Detects Differences in Recovery after Spinal Cord Injury in Five Common Mouse Strains. Journal of Neurotrauma 23, 635–659 (2006).

41. Brennan, F. H. et al. Microglia coordinate cellular interactions during spinal cord repair in mice. Nat Commun 13, 4096 (2022).

42. Milich, L. M. et al. Single-cell analysis of the cellular heterogeneity and interactions in the injured mouse spinal cord. Journal of Experimental Medicine 218, e20210040 (2021).

43. Wang, J. et al. Single-cell transcriptome analysis reveals the immune heterogeneity and the repopulation of microglia by Hif1α in mice after spinal cord injury. Cell Death Dis 13, 432 (2022).

44. Grieshaber-Bouyer, R. et al. The neutrotime transcriptional signature defines a single continuum of neutrophils across biological compartments. Nat Commun 12, 2856 (2021).

45. Wigerblad, G. et al. Single-Cell Analysis Reveals the Range of Transcriptional States of Circulating Human Neutrophils. J Immunol 209, 772–782 (2022).

46. Aran, D. et al. Reference-based analysis of lung single-cell sequencing reveals a transitional profibrotic macrophage. Nat Immunol 20, 163–172 (2019).

47. Zhou, Y. et al. Metascape provides a biologist-oriented resource for the analysis of systems-level datasets. Nat Commun 10, 1523 (2019).

48. Vafadarnejad, E. et al. Dynamics of Cardiac Neutrophil Diversity in Murine Myocardial Infarction. Circ Res 127, e232–e249 (2020).

49. Boivin, G. et al. Durable and controlled depletion of neutrophils in mice. Nat Commun 11, 2762 (2020).

50. Soehnlein, O. & Lindbom, L. Phagocyte partnership during the onset and resolution of inflammation. Nat Rev Immunol 10, 427–439 (2010).

51. Stewart, A. N. et al. Nonresolving Neuroinflammation Regulates Axon Regeneration in Chronic Spinal Cord Injury. J. Neurosci. 45, e1017242024 (2025).

52. Castanheira, F. V. S. & Kubes, P. Neutrophils and NETs in modulating acute and chronic inflammation. Blood 133, 2178–2185 (2019).

53. Soehnlein, O., Steffens, S., Hidalgo, A. & Weber, C. Neutrophils as protagonists and targets in chronic inflammation. Nat Rev Immunol 17, 248–261 (2017).

54. Taoka, Y. et al. Role of neutrophils in spinal cord injury in the rat. Neuroscience 79, 1177–1182 (1997).

55. Saiwai, H. et al. The LTB4-BLT1 Axis Mediates Neutrophil Infiltration and Secondary Injury in Experimental Spinal Cord Injury. The American Journal of Pathology 176, 2352–2366 (2010).

56. Devi, S. et al. Neutrophil mobilization via plerixafor-mediated CXCR4 inhibition arises from lung demargination and blockade of neutrophil homing to the bone marrow. J Exp Med 210, 2321–2336 (2013).

57. Jogia, T. et al. Prognostic value of early leukocyte fluctuations for recovery from traumatic spinal cord injury. Clinical and Translational Medicine 11, e272 (2021).

58. Herisson, F. et al. Direct vascular channels connect skull bone marrow and the brain surface enabling myeloid cell migration. Nat Neurosci 21, 1209–1217 (2018).

59. Ng, M. S. F. et al. Deterministic reprogramming of neutrophils within tumors. Science 383, eadf6493 (2024).

60. Ballesteros, I. et al. Co-option of Neutrophil Fates by Tissue Environments. Cell 183, 1282–1297.e18 (2020).

61. Hahn, J. et al. Aggregated neutrophil extracellular traps resolve inflammation by proteolysis of cytokines and chemokines and protection from antiproteases. FASEB J 33, 1401–1414 (2019).

62. Cauwe, B., Martens, E., Proost, P. & Opdenakker, G. Multidimensional degradomics identifies systemic autoantigens and intracellular matrix proteins as novel gelatinase B/MMP-9 substrates. Integr Biol (Camb) 1, 404–426 (2009).

63. Bosurgi, L. et al. Macrophage function in tissue repair and remodeling requires IL-4 or IL-13 with apoptotic cells. Science 356, 1072–1076 (2017).

