## Supplemental Figures for "Mature neutrophils promote long-term functional recovery after spinal cord injury in a sex-dependent manner"

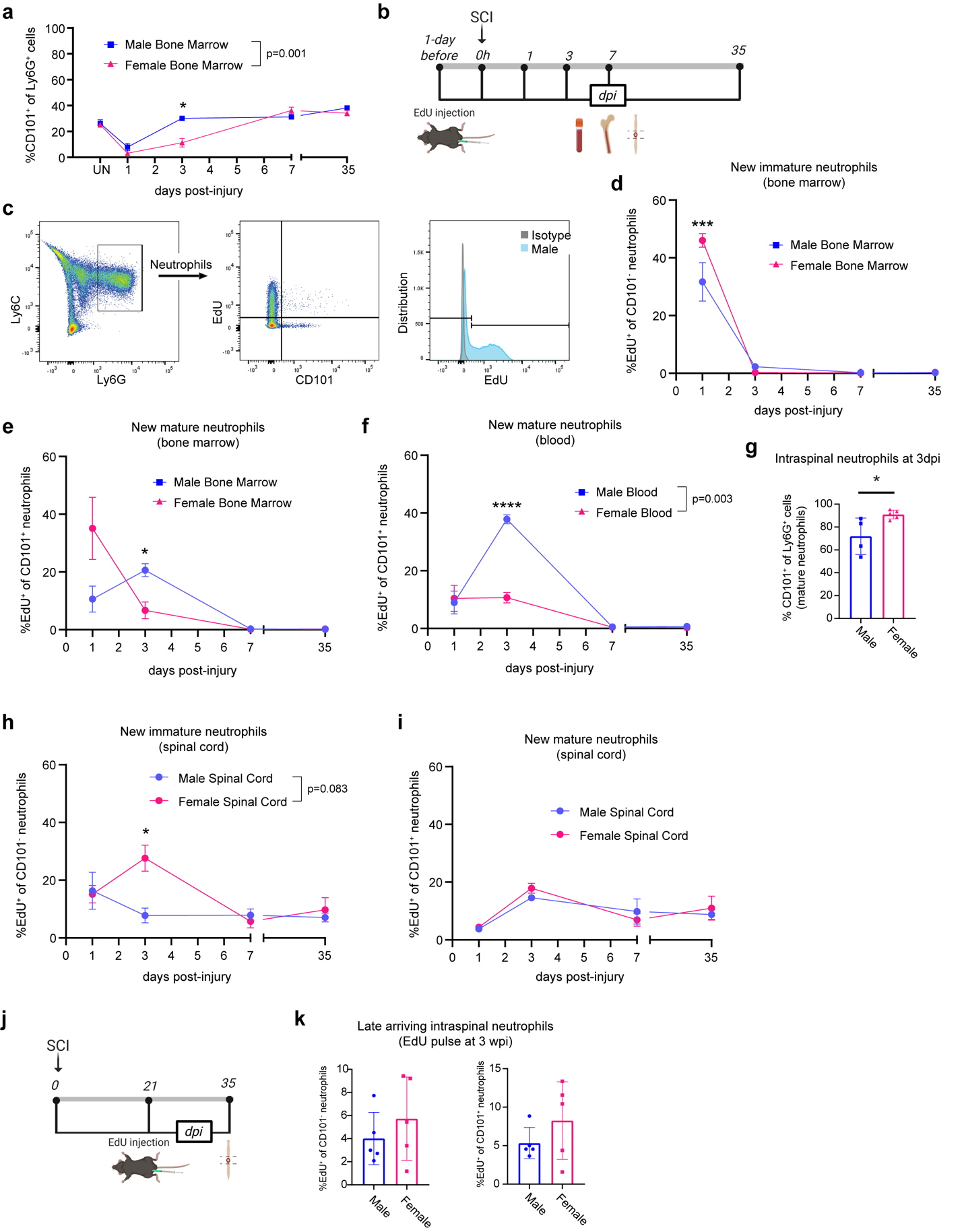

**Supplemental Figure 1: Systemic and localized neutrophil populations change after SCI in a sex-dependent manner.** (a) Quantification of the percentage of mature (CD101^+^) neutrophils out of the total neutrophil population (Ly6G^+^ cells) by flow cytometry in the bone marrow after SCI. The proportion of mature neutrophils is greater in male mice vs. females at 3dpi. n=3-6/sex/treatment. Mixed effects analysis with Sidak’s multiple comparisons test. (b) Schematic of EdU injection at 1 day prior to SCI and tissue collection for flow cytometry analysis at 1, 3, 7 and 35 dpi. (c) Representative flow cytometry gating of neutrophils and EdU^+^ neutrophils. (d) Quantification of EdU^+^ immature neutrophils (CD101^-^/Ly6G^+^ cells) in the bone marrow after SCI. n=5-6/sex/treatment. Two-way ANOVA with Sidak’s multiple comparisons test. (e) Quantification of EdU^+^ mature neutrophils (CD101^+^/Ly6G^+^ cells) in the bone marrow after SCI. n=5-6/sex/treatment. Mixed effects analysis with Sidak’s multiple comparisons test. (f) Quantification of EdU^+^ mature neutrophils (CD101^+^/Ly6G^+^ cells) in the blood after SCI. n=5-6/sex/treatment. Mixed effects analysis with Sidak’s multiple comparisons test. (g) Quantification of mature neutrophils (CD101^+^/Ly6G^+^ cells) in the spinal cord at 3 dpi. n=4-5/sex/treatment. Unpaired two-tailed Student’s t-test. (h) Quantification of EdU^+^ immature neutrophils (CD101^-^/Ly6G^+^ cells) in the spinal cord after SCI. n=5-6/sex/treatment. Mixed effects analysis with Sidak’s multiple comparisons test. (i) Quantification of EdU^+^ mature neutrophils (CD101^+^/Ly6G^+^ cells) in the spinal cord after SCI. n=5-6/sex/treatment. Mixed effects analysis with Sidak’s multiple comparisons test. (j) Schematic of EdU injection at 21 dpi and tissue collection for flow cytometry analysis at 35 dpi. (k) Quantification of EdU^+^ immature neutrophils (CD101^-^/Ly6G^+^ cells; left) and mature neutrophils (CD101^+^/Ly6G^+^ cells, right) in the spinal cord after SCI. n=5/sex/treatment. Unpaired two-tailed Student’s t-test. *p<0.05, ****p<0.0001. Abbreviations: SCI = spinal cord injury, dpi = days post-injury. Mean ± SEM. wpi = weeks post-injury

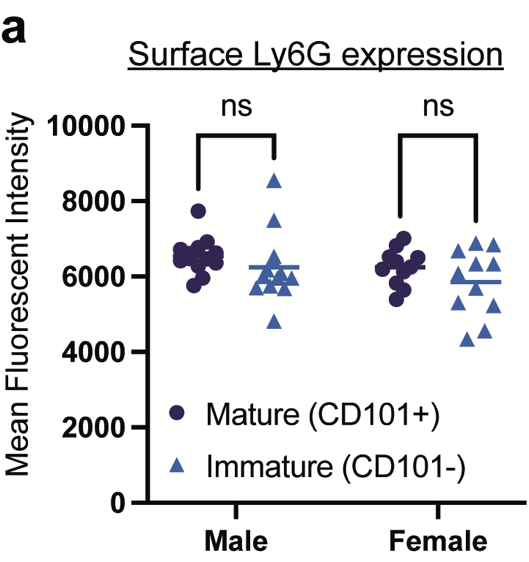

**Supplemental Figure 2: Ly6G levels on immature and mature neutrophils.** (a) Mean fluorescent intensity of surface Ly6G labeling on mature (CD101^+^) and immature (CD101^-^) neutrophils in the blood of uninjured mice as assessed via flow cytometry. No difference was observed for surface Ly6G levels. n =11-13/sex/treatment. Two-way ANOVA with Sidak’s multiple comparisons post-hoc test. **p<0.01. Mean ± SEM.

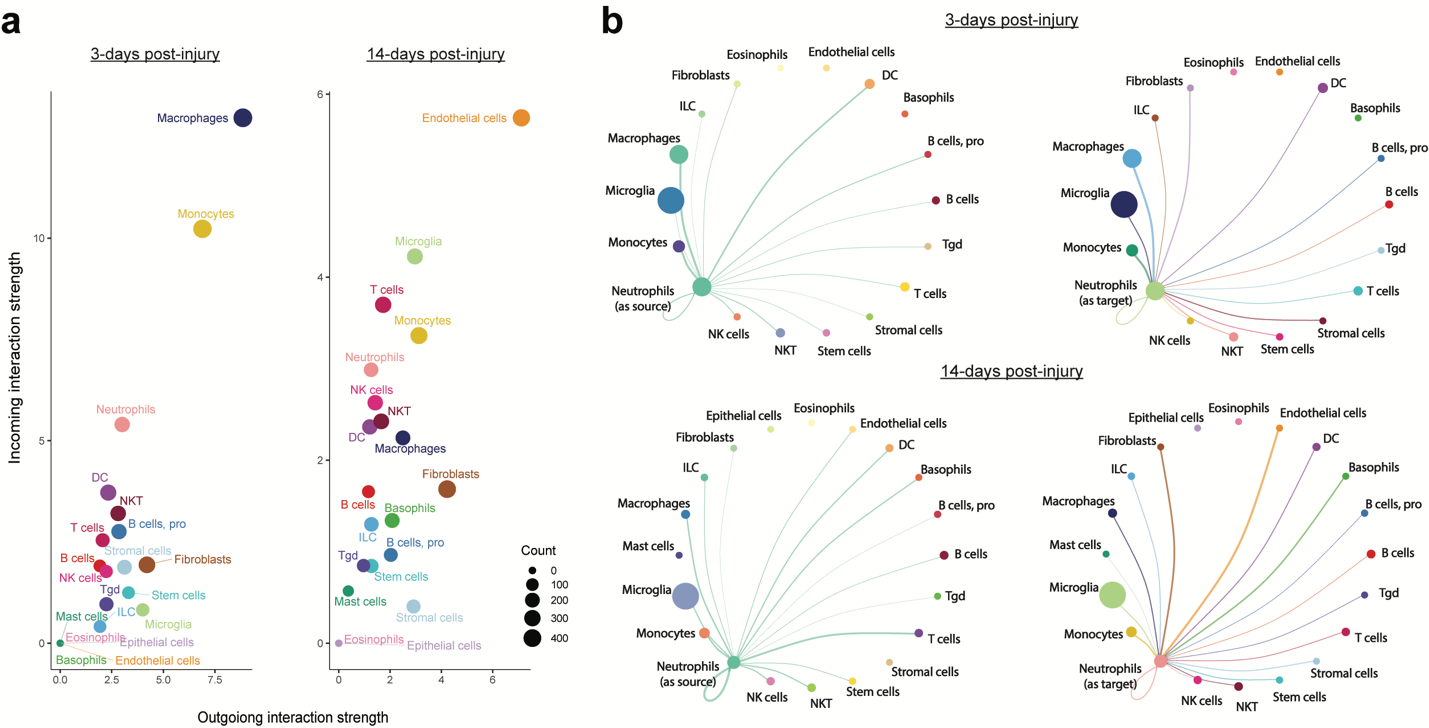

**Supplemental Figure 3: CellChat analysis of intraspinal neutrophil receptor-ligand interactions.** (a) Incoming and outgoing receptor-ligand interaction strengths for intraspinal neutrophils at 3 and 14 dpi from Wang *et al*. 2022. Neutrophils are predicted to interact strongly with macrophages at 3 dpi. (b) Outgoing (left) and incoming (right) interaction maps for intraspinal neutrophils with other CD45^+^ cells at 3 and 14 dpi.

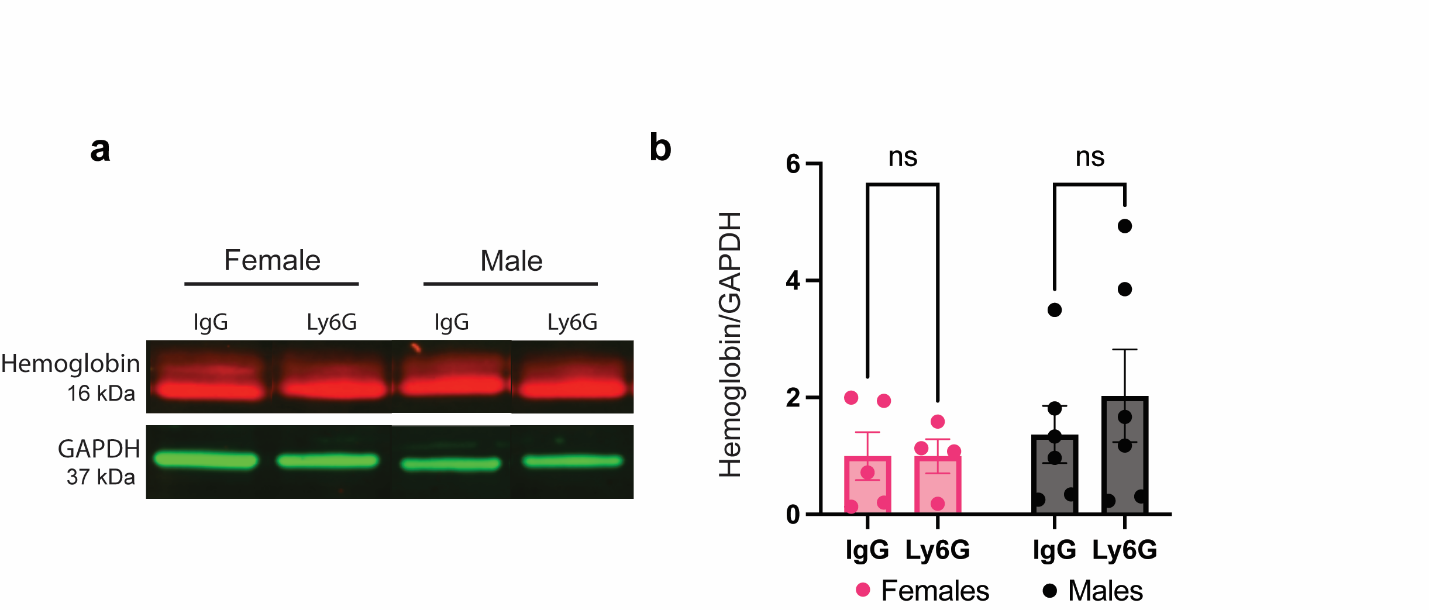

**Supplemental Figure 4: Neutrophil depletion does not alter acute blood-spinal cord barrier disruption.** (a) Representative western blot analysis of hemoglobin (as a surrogate for red blood cells) in the injured spinal cord at 1 dpi. (b) Quantification of hemoglobin levels normalized to GAPDH. n=4-6/sex/treatment. Two-way ANOVA with Sidak’s multiple comparisons post-hoc test. Mean ± SEM.

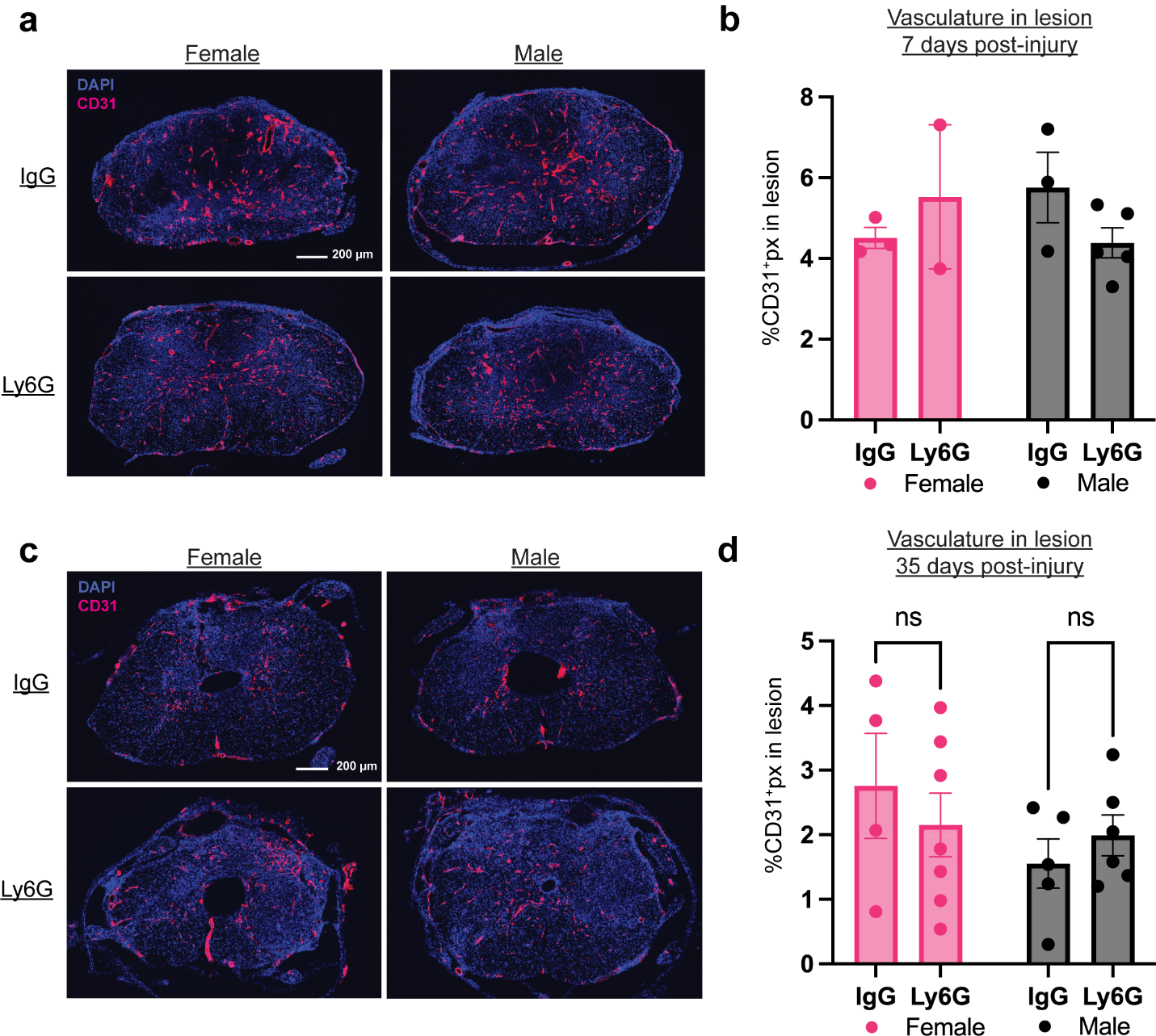

**Supplemental Figure 5: Neutrophil depletion does not alter vasculature damage in the subacute and subchronic phases of SCI.** (a, c) Representative images showing CD31 (pink) and DAPI (blue) labeling in male and female mice with neutrophil depletion (Ly6G) or control (IgG) antibody at 7 (a) and 35 (c) dpi. (b) Quantification of CD31 immunolabeling in the injured spinal cord at 7 dpi from five serial sections spanning 2.25 mm centered on the lesion. n=2-5/sex/treatment. (d) Quantification of CD31 (percentage of total pixels that were CD31^+^) immunolabeling in the injured spinal cord at 35 dpi from five serial sections spanning 1.25 mm centered on the lesion. n=4-7/sex/treatment. Two-way ANOVA with Sidak’s multiple comparisons post-hoc test. Mean ± SEM.

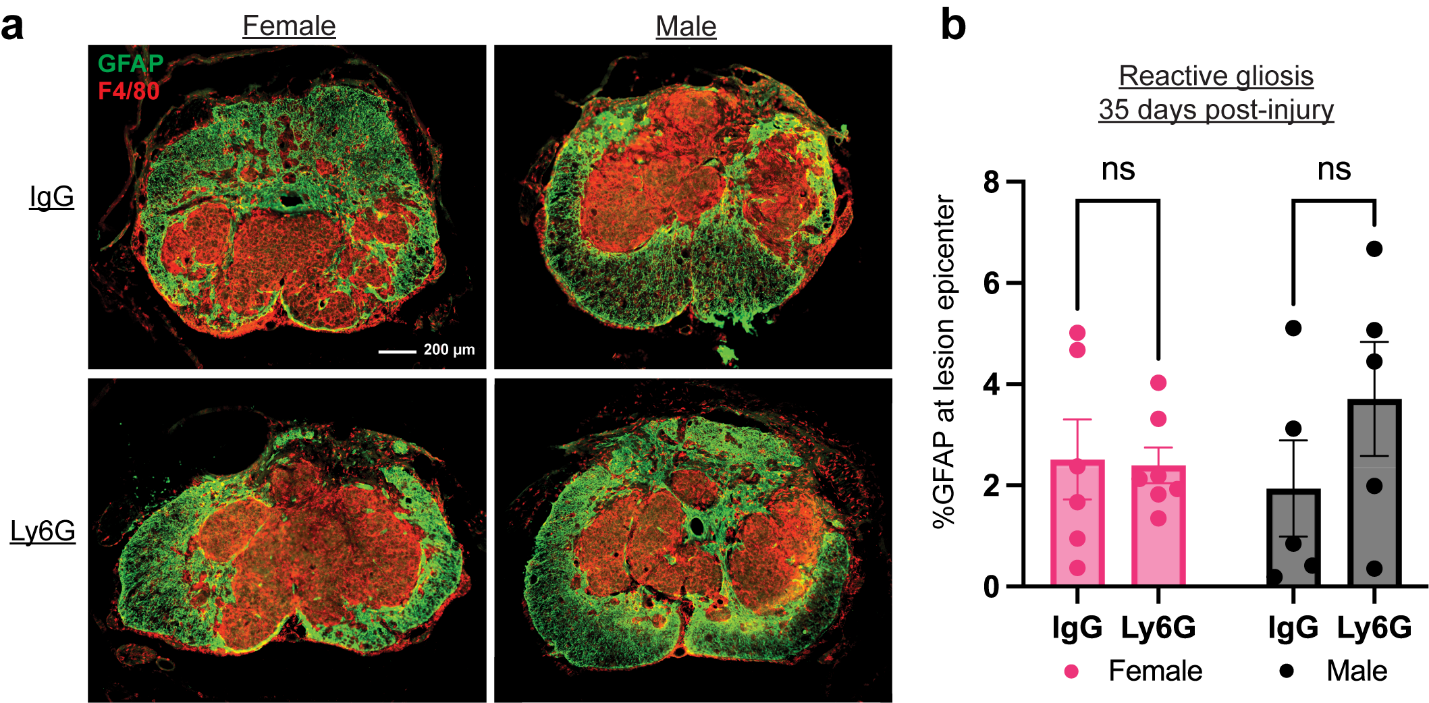

**Supplemental Figure 6: Neutrophil depletion does not alter glial scar formation after SCI.** (a) Representative images showing GFAP (green) for reactive gliosis and F4/80 (red) for macrophages at the lesion epicenter in male and female mice with neutrophil depletion (Ly6G) or control (IgG) antibody. (b) Quantification of GFAP immunolabeling in the injured spinal cord at 35 dpi from five serial sections spanning 1mm centered on the lesion epicenter. n=5-7/sex/treatment. Two-way ANOVA with Sidak’s multiple comparisons post-hoc test. Mean ± SEM.
